# Instant fluorescence lifetime imaging microscopy reveals mechano-metabolic reprogramming of stromal cells in breast cancer peritumoral microenvironments

**DOI:** 10.1101/2025.05.28.656717

**Authors:** Julian Najera, Hao Chen, Bianca Batista, Frank Ketchum, Aktar Ali, Pinar Zorlutuna, Scott Howard, Meenal Datta

**Affiliations:** Department of Aerospace and Mechanical Engineering, University of Notre Dame, Notre Dame, IN, USA.; Bioengineering Graduate Program, University of Notre Dame, Notre Dame, IN, USA.; Department of Electrical Engineering, University of Notre Dame, Notre Dame, IN, USA.; Department of Chemical and Biomolecular Engineering, University of Notre Dame, Notre Dame, IN, USA.; Department of Chemistry and Biochemistry, University of Notre Dame, Notre Dame, IN, USA.

**Keywords:** solid stress, host tissue, glycolysis, oxidative phosphorylation, mitochondria

## Abstract

The breast peritumor microenvironment (pTME) is increasingly recognized as a mediator of breast cancer progression and treatment resistance. However, if and how growth-induced tumor compressive forces (i.e., solid stresses) influence the breast pTME remains unclear. Here we show using instant fluorescence lifetime imaging microscopy (FLIM)—a frequency-domain FLIM system capable of simultaneous image acquisition and instantaneous data processing—that breast tumor-mimicking *in vitro* compression promotes metabolic changes in stromal cells found in the breast pTME. Namely, compression shifts NIH3T3 fibroblasts and differentiated 3T3-L1 (d3T3-L1) adipocytes toward a more glycolytic state, while it promotes increased oxidative phosphorylation in 3T3-L1 undifferentiated adipocytes. The gold-standard Seahorse extracellular flux assay fails to capture these changes, underscoring the superior sensitivity of instant FLIM in detecting metabolic shifts. We validate these phenotypic findings at the transcriptomic level via RNA sequencing, confirming that compressed fibroblasts downregulate oxidative phosphorylation and upregulate glycolysis compared to uncompressed controls. We further demonstrate that compression induces mitochondrial dysregulation in undifferentiated adipocytes, driven in part by upregulated mitophagy and disrupted fusion dynamics. Finally, we confirm that these stromal cell types recapitulate these distinct metabolic states in human breast cancer patient samples, consistent with our *in vitro* findings. By elucidating mechano-metabolic interactions occurring at the tumor-host interface, these results will inform the development of innovative mechano-metabolic reprogramming treatment strategies to improve breast cancer patient survival.

## Introduction

Despite advances in detection and therapy, breast cancer remains the most frequently diagnosed cancer and leading cause of cancer-related death in women globally.^1-3^ Once considered healthy, the breast peritumor—non-malignant tissue adjacent to the tumor mass—is now recognized to harbor tumor-supportive properties.^4-9^ Peritumoral adipocytes and fibroblasts, for instance, are metabolically reprogrammed to produce essential metabolites (e.g., amino acids, nutrients, fatty acids, and ketone bodies) for nearby breast cancer cells to exploit to meet their own bioenergetic needs.^10,11^ Notably, this metabolic coupling has been linked to drug resistance,^12,13^ suggesting that treatment response can be improved by targeting the altered metabolic landscape of the breast peritumor microenvironment (pTME).

While these metabolic insights have largely focused on biochemical drivers, emerging evidence suggests that aberrant physical cues may also influence peritumoral stromal cell metabolism. It is becoming increasingly clear that mechanical *properties* (e.g., elevated stiffness) regulate the metabolism of both malignant and non-malignant cells in the tumor during different stages of disease progression.^14-17^ However, the extent to which mechanical *forces*—particularly the compressive and deformative solid stresses exerted by the mammary tumor onto the surrounding host tissue during growth and expansion^18,19^—impact the metabolic state of stromal cells in the breast pTME remains unknown.

Metabolic cell profiling techniques have rapidly evolved in recent years, including photonics-based methods such as fluorescence lifetime imaging microscopy (FLIM). FLIM is an advanced technique that distinguishes overlapping emission spectra based on differences in the fluorescence lifetimes of fluorophores, including cell-endogenous fluorophores.^20,21^ By measuring the autofluorescence of the metabolic co-factor NADH, FLIM provides a non-destructive, label-free approach to detect cellular metabolic changes. This is because NADH exhibits longer lifetimes (𝜏) in the protein-bound state and shorter lifetimes in the free state.^22^ Therefore, shorter mean lifetimes (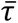) correlate with enhanced glycolysis, while higher 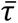 is associated with increased oxidative phosphorylation (OXPHOS).

The fluorescence lifetime of a fluorophore can be measured in the time-domain (TD) or frequency-domain (FD), with each technique offering a unique set of challenges.^20^ Time-correlated single-photon counting (TCSPC)—the most widely used and commercially available TD-FLIM technique—is often limited by slow data processing times, hardware cost, data volume, and large computer memory requirements.^20,23^ On the other hand, FD-FLIM is typically limited by its low signal-to-noise ratio (SNR) in lifetime measurements and relatively difficult implementation. To address both sets of challenges, we recently developed a novel, cost- and time-effective FD-FLIM technique known as “instant FLIM.”^23^ Using time-domain data acquisition coupled with an analog signaling processing method in FD, instant FLIM is capable of simultaneously collecting high-resolution images and performing instantaneous data processing for lifetime quantification. Importantly, we previously demonstrated that the SNR performance of instant FLIM is superior to that of conventional FD-FLIM which typically implement a detector gain modulator as the modulation source.^23^

Here, we sought to elucidate how compressive solid stress influences breast pTME stromal metabolism and demonstrate the application of instant FLIM as a practical and effective platform to address this gap in the field. Using an *in vitro* compression device (**Fig. 1a**) with instant FLIM (**Fig. 1b**), we discovered that breast tumor-mimicking compression drives a glycolytic shift in NIH3T3 fibroblasts and differentiated 3T3-L1 (d3T3-L1) adipocytes, and an oxidative shift in 3T3-L1 undifferentiated adipocytes. These changes were not detected by less sensitive metabolic assays (e.g., Seahorse extracellular flux [XF]), but were confirmed by bulk-RNA sequencing which showed that compressed fibroblasts undergo significant transcriptional changes indicative of glycolytic reprogramming. We further discovered that compression induces mitochondrial fragmentation in undifferentiated adipocytes by impairing fusion dynamics, potentially as a consequence of increased mitophagy. Interestingly, the introduction of 4T1 murine triple-negative breast cancer cell-conditioned media (CM) abrogates compression-induced metabolic reprogramming in fibroblasts and undifferentiated adipocytes but not in differentiated adipocytes. Finally, our analysis of retrospective data from human breast cancer tissue samples and single cell RNA sequencing (scRNA-seq) data revealed spatially-dependent metabolic reprogramming of peritumoral fibroblasts and adipocytes consistent with our *in vitro* findings. Collectively, these findings provide new insight into tumor-host mechano-metabolic crosstalk with translational significance.

**Figure 1.**
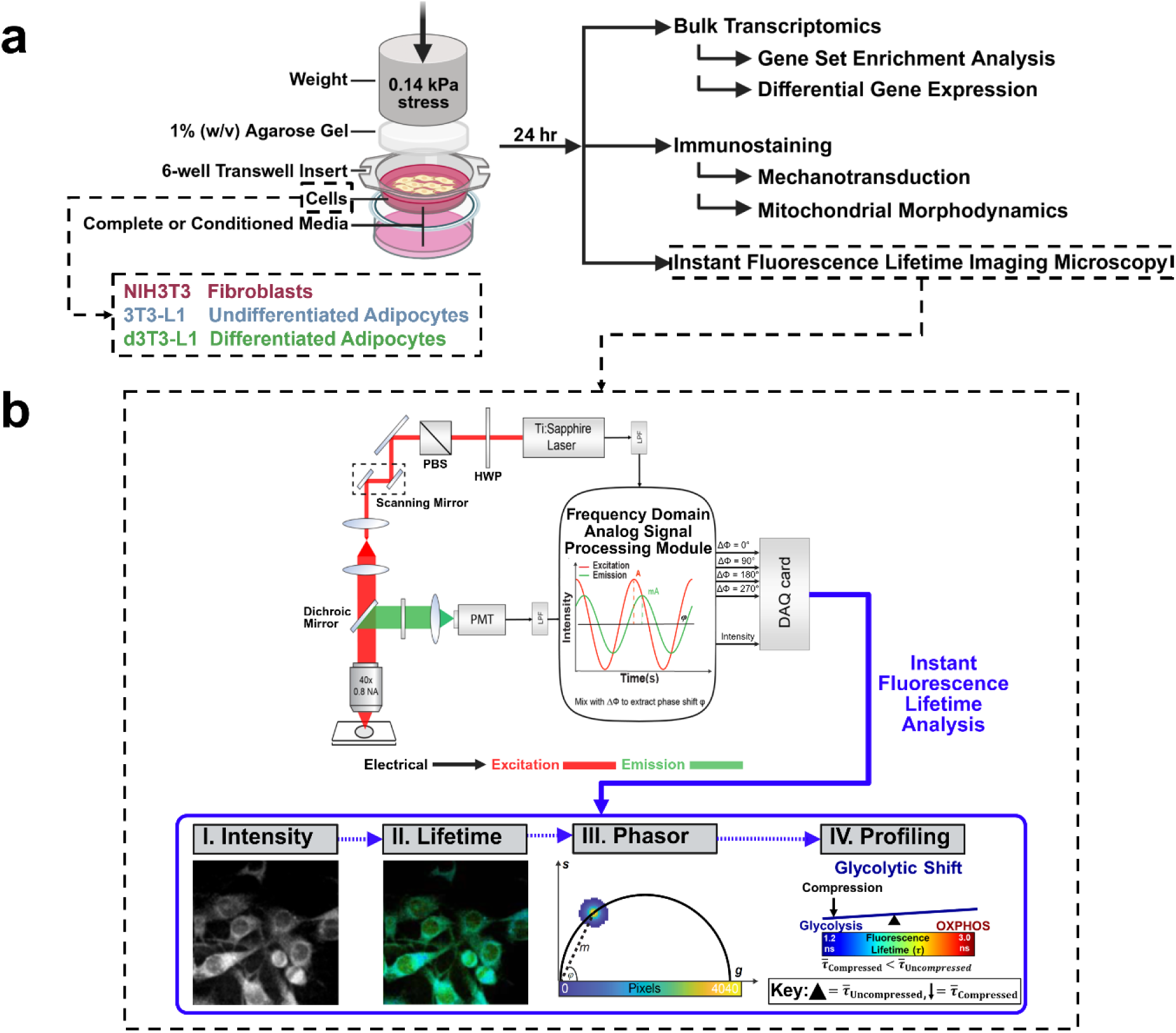
Schematic of experimental approach and instant FD-FLIM system. Stromal cells found in the breast peritumor experience breast tumor-mimicking compression^79^ (∼0.14 kPa) for 24 hours using an **a)** *in vitro* compression apparatus. The resulting response is assessed using omics, immunofluorescence, and **b)** instant fluorescence lifetime imaging microscopy (FLIM). In instant FLIM, a femtosecond Ti:Sapphire laser generates an excitation light (red). The excited fluorescence signal (green) enters a photomultiplier tube (PMT), is converted into an electrical signal, then enters an analog signal processing module. Through homodyne detection, the signal is mixed with multiple phase-shifted components from the excitation signal to simultaneously generate intensity and four phase-shifted mixed signals. This data is digitized by a data acquisition (DAQ) card (**b-I,** blue), then processed and displayed as a lifetime image and phasor plot (**b-II** and **b-III**). Phasor plots, defined by g and s components, visualize fluorescence lifetimes (𝜏), wherein points closer to the g-axis correspond to shorter lifetimes. The distance from the origin (m) reflects the modulation depth, while the angle (𝜑) represents lifetime dynamics (**b-III**). The color bar represents lifetime values coded from lowest (blue) to highest (red). Metabolic changes are assessed by comparing the overall mean lifetime (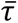) of compressed cells (downward arrow) to uncompressed ones (fulcrum; **b-IV**). Relative to the fulcrum, lower 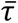 indicates a glycolytic shift (shown here) while higher 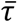 reflects an oxidative shift. HWP: half-wave plate, PBS: polarizing beam splitter, LPF: low-pass filter, A: ampere, mA: milliampere. Image in **a)** created with Biorender.com., https://biorender.com/.

## Results

### Implementation, principles, and application of instant FLIM to pTME samples

Based on our previous work,^23^ instant FLIM combines the advantages of time-domain data acquisition and sampling to provide real-time fluorescence intensity decay measurements with high sensitivity. This significantly improves SNR, reduces data loss, and enables faster, more efficient analysis without requiring extra computation time or memory.

Instant FLIM enables non-destructive observation of small changes in cell metabolism with deep penetration and high resolution, providing a consistent and reliable approach for studying metabolic responses in the dynamic microenvironment of breast cancer. By measuring intrinsic autofluorescence in cells, our system is capable of detecting lifetime changes with precision on the order of 100 picoseconds compared to the nanosecond-level lifetimes typically observed. Specifically, for breast cancer metabolism microenvironment dynamics, the glycolysis pathway can be reflected by NAD(P)H and its binding with proteins. With estimated NAD(P)H lifetime ranges, we can precisely detect metabolic shifts at the subcellular level, as described in detail in the results below.

Specifically, when examining complex and dynamic biological microenvironments in *in vitro* samples (as described in subsequent sections), fluorescence lifetime imaging typically reveals heterogeneous populations of fluorophores, with further distortion caused by formalin fixation.^24-26^ As such, instant FLIM improves the stability of measurement results through rapid averaging and by applying the phasor approach to segment and quantify the contributions of different fluorophores based on their relative contributions to the measurement (**Fig. 1b**), thereby minimizing the effects of background signals. Additionally, we demonstrate in our final results that instant FLIM can strongly differentiate adipocytes associated with different regions of interest within and near the pTME.

### Instant FLIM reveals altered metabolic profiles in compressed fibroblasts and adipocytes

We utilized a previously established transwell compression system^27-29^ (**Fig. 1a**) coupled with instant FLIM (**Fig. 1b**) to assess the effect of 24 hours of *in vitro* compression (∼0.14 kPa) on the metabolic state of NIH3T3 fibroblasts (**Fig. 2a**), 3T3-L1 undifferentiated adipocytes (**Fig. 2b**), and d3T3-L1 differentiated adipocytes (**Fig. 2c**). Using phasor-based fluorescence lifetime analysis, we determined that the application of compressive stress significantly alters fluorescence lifetime in all three cell types, thereby indicating a change in cell metabolism. Response to mechanical stress appears to be heterogenous among different cell types, however, as the lifetime in compressed fibroblasts (**Fig. 2d**) and differentiated adipocytes decreases (**Fig. 2f**), but increases in compressed undifferentiated adipocytes (**Fig. 2e**). These results suggest that compression metabolically reprograms fibroblasts and adipocytes toward a more glycolytic state (**Fig. 2g,i**), while enhancing oxidative phosphorylation in undifferentiated adipocytes (**Fig. 2h**).

**Figure 2.**
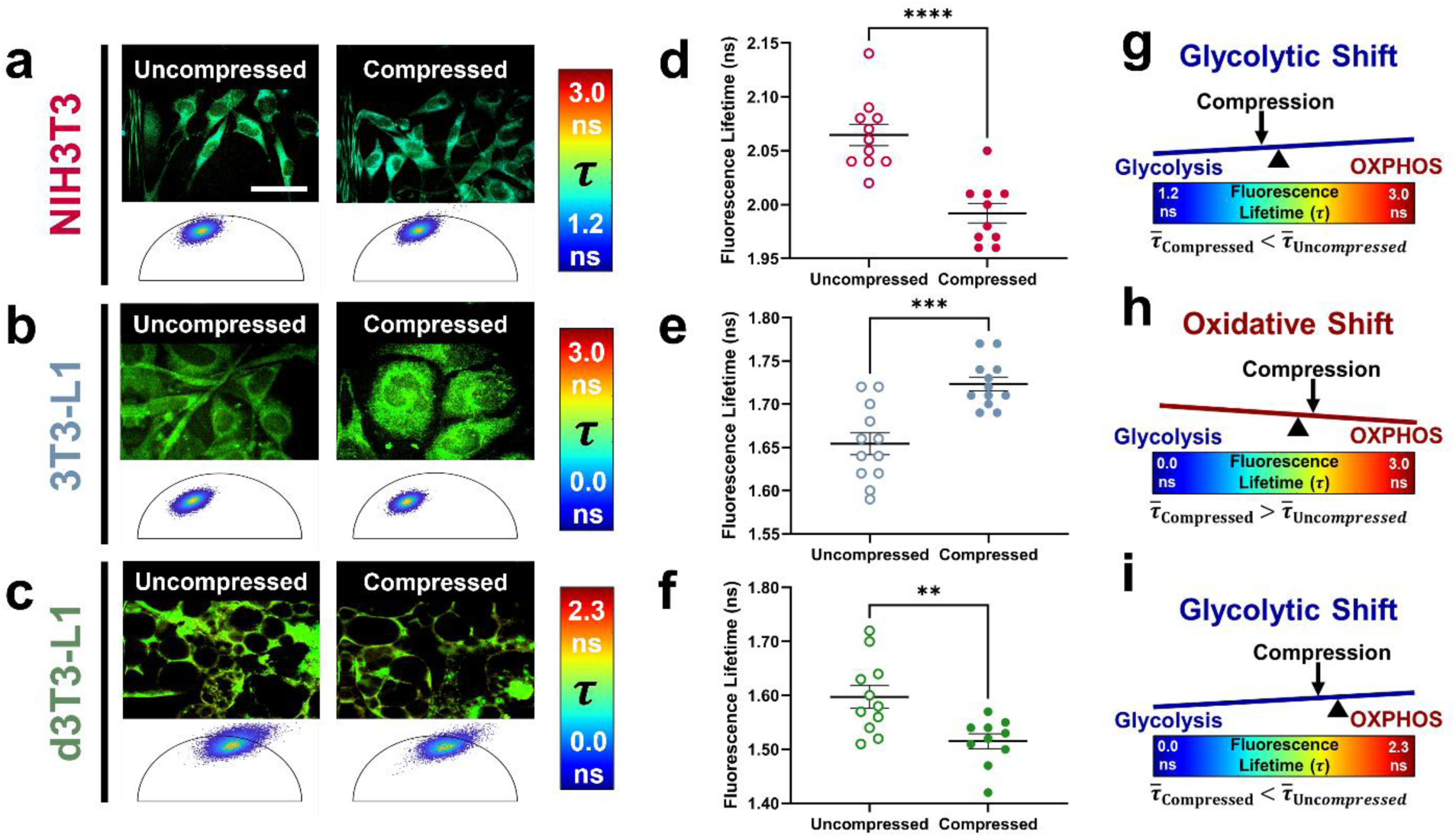
Breast tumor-mimicking compression promotes differential metabolic shifts in peritumoral stromal cell types. Representative fluorescence lifetime images (top) and phasor plots (bottom) of **a)** NIH3T3 fibroblasts (n = 11 uncompressed, 10 compressed), **b)** 3T3-L1 undifferentiated adipocytes (n = 12 uncompressed and compressed), and **c)** differentiated 3T3-L1 (d3T3-L1) adipocytes (n = 11 uncompressed, 10 compressed). Fluorescence lifetime analysis reveals that the applied mechanical stress decreases mean fluorescence lifetime (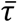) in **d)** fibroblasts and **f)** differentiated adipocytes, but produces the opposite effect in **e)** undifferentiated adipocytes. The decrease in lifetime indicates that compression metabolically rewires **g)** NIH3T3 and **i)** d3T3-L1 cells to a more glycolytic state and **h)** 3T3-L1s towards a more oxidative state. Error bars represent SEM and asterisks indicate statistical significance (\**p* < 0.05, \*\**p* < 0.01, \*\*\**p* < 0.001, \*\*\*\**p* < 0.0001) determined using a Student’s t-test. Scale bar is 50 µm.

### In vitro compression activates YAP in fibroblasts and undifferentiated adipocytes

To determine whether *in vitro* compression triggers a mechanical response, we evaluated the transcription of key mechanotransducers (*Ctnnb1*, *Piezo1*, *Trpv4*, and *Yap1*) using qPCR in NIH3T3 and 3T3-L1 cells. We found that applied compression does not promote significant differences in the expression of *Ctnnb1*, *Piezo1,* and *Yap1* in either cell type, while *Trpv4* is moderately upregulated in compressed fibroblasts, but not undifferentiated adipocytes (**Fig. S1**).

Given that the transcriptional co-factor, YAP, is known to translocate to the nucleus in response to mechanical cues,^30^ however, we next analyzed the cellular localization of the protein. By examining the intensity ratio of nuclear YAP to perinuclear YAP (nuclear:perinuclear intensity) in immunofluorescently-stained samples (**Fig. 3a,b**), we discovered that compressed NIH3T3 and 3T3-L1 cells exhibit higher levels of nuclear YAP compared to their uncompressed counterparts (**Fig. 3c,d**). Overall, our results confirm that both fibroblasts and undifferentiated adipocytes exhibit a mechanobiological response to compressive stress *in vitro* by increasing YAP activation.

**Figure 3.**
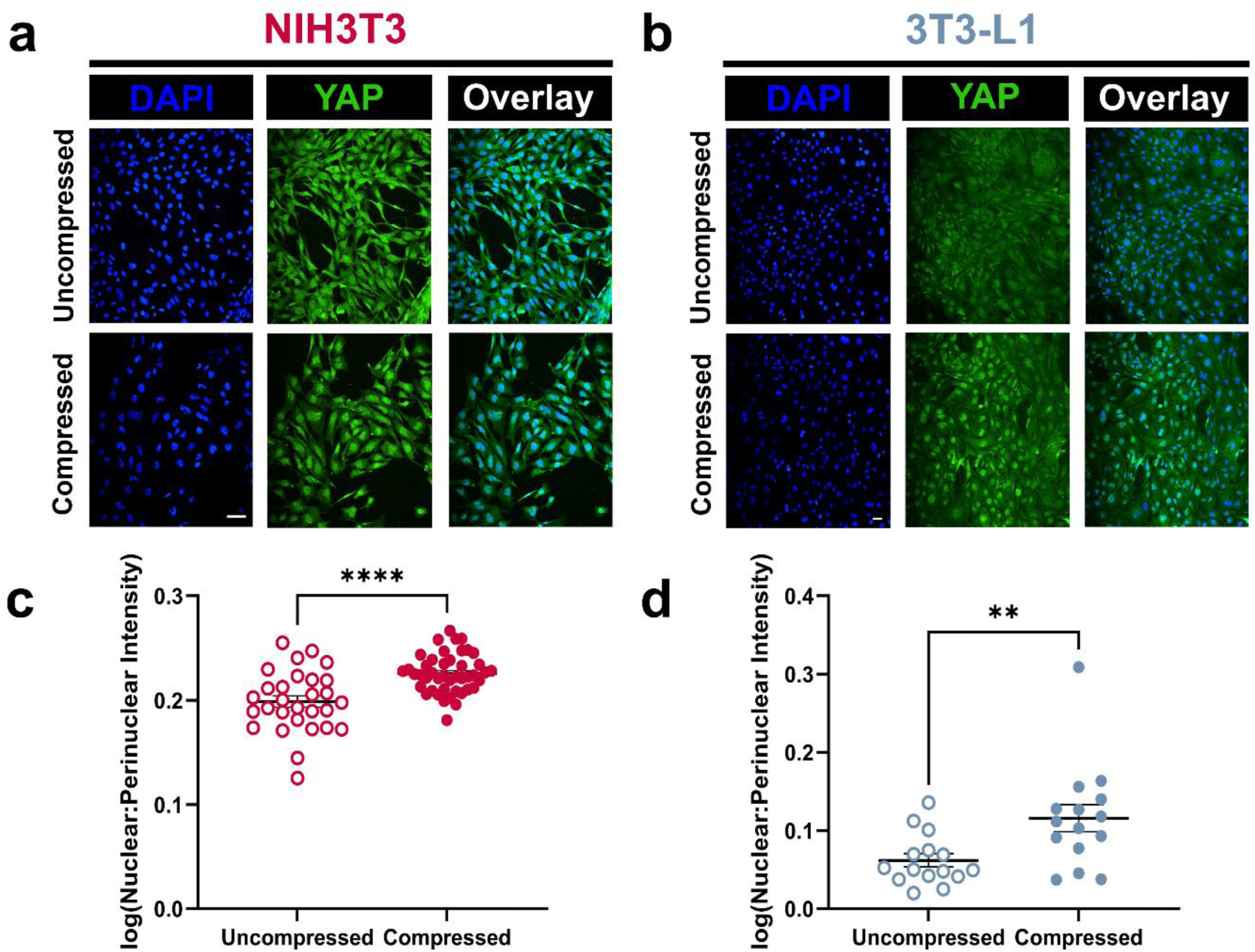
Compression promotes YAP activation in fibroblasts and undifferentiated adipocytes. Representative immunofluorescent images of murine **a)** NIH3T3 (n = 29 uncompressed, 44 compressed) and **b)** 3T3-L1 cells (n = 15 uncompressed and compressed) stained for the mechanoresponsive yes-associated protein (YAP). Analysis of log-transformed nuclear-to-perinuclear YAP intensity ratios (log[nuclear:perinuclear intensity]) reveals that compression significantly increases YAP nuclear localization in **c)** fibroblasts and **d)** undifferentiated adipocytes. Error bars represent SEM and asterisks indicate statistical significance (\**p* < 0.05, \*\**p* < 0.01, \*\*\**p* < 0.001, \*\*\*\**p* < 0.0001) determined using a Student’s t-test. Scale bar is 50 µm.

### Instant FLIM offers superior sensitivity over real-time ATP analysis in capturing metabolic changes under compression

The Seahorse XF Real-Time ATP Rate Assay is commonly used to evaluate changes in energy metabolism in live cells by measuring and quantifying ATP production rates via glycolysis and mitochondrial respiration in real time. To validate our FLIM findings, we performed this assay under the same compression conditions (**Fig. S3a**). Interestingly, we did not observe significant differences in ATP production from either metabolic pathway between compressed and uncompressed cells (**Fig. S3b,c**). Furthermore, we did not identify an increased reliance on energy production from one pathway over the other (**Fig. S3d**). This suggests that instant FLIM provides enhanced sensitivity in detecting compression-induced metabolic alterations in cells compared to conventional platforms such as Seahorse.

### Bulk-RNA sequencing analysis confirms metabolic reprogramming at the transcriptional level in compressed fibroblasts

We next conducted functional enrichment analysis of bulk-RNA sequencing data collected from NIH3T3 cells to confirm that compression-induced metabolic reprogramming occurs at the transcriptional level. Our analysis revealed that compression significantly downregulates genes associated with OXPHOS and related processes, while concurrently promoting the upregulation of transcripts involved in glucose metabolism, glucose import, and the glycolytic process to varying degrees of significance (**Fig. 4a,b**). For reference, **Supplemental Table 1** contains the full list of glycolysis- and OXPHOS-related transcripts and their corresponding log2 fold-changes. Additionally, a list of the top differentially expressed genes is provided in **Supplemental Table 2**. Complementing our transcriptomic findings, a glucose uptake assay showed that compression increases glucose uptake in fibroblasts (**Fig. S2**). Together, these results indicate that, at the transcriptional and functional levels, compression drives a glycolytic transition in fibroblast metabolism, with reduced reliance on OXPHOS. Additionally, it underscores the advantage of instant FLIM as being a system capable of capturing subtle yet significant metabolic differences *in situ* that may have otherwise been overlooked by traditional functional assays.

**Figure 4.**
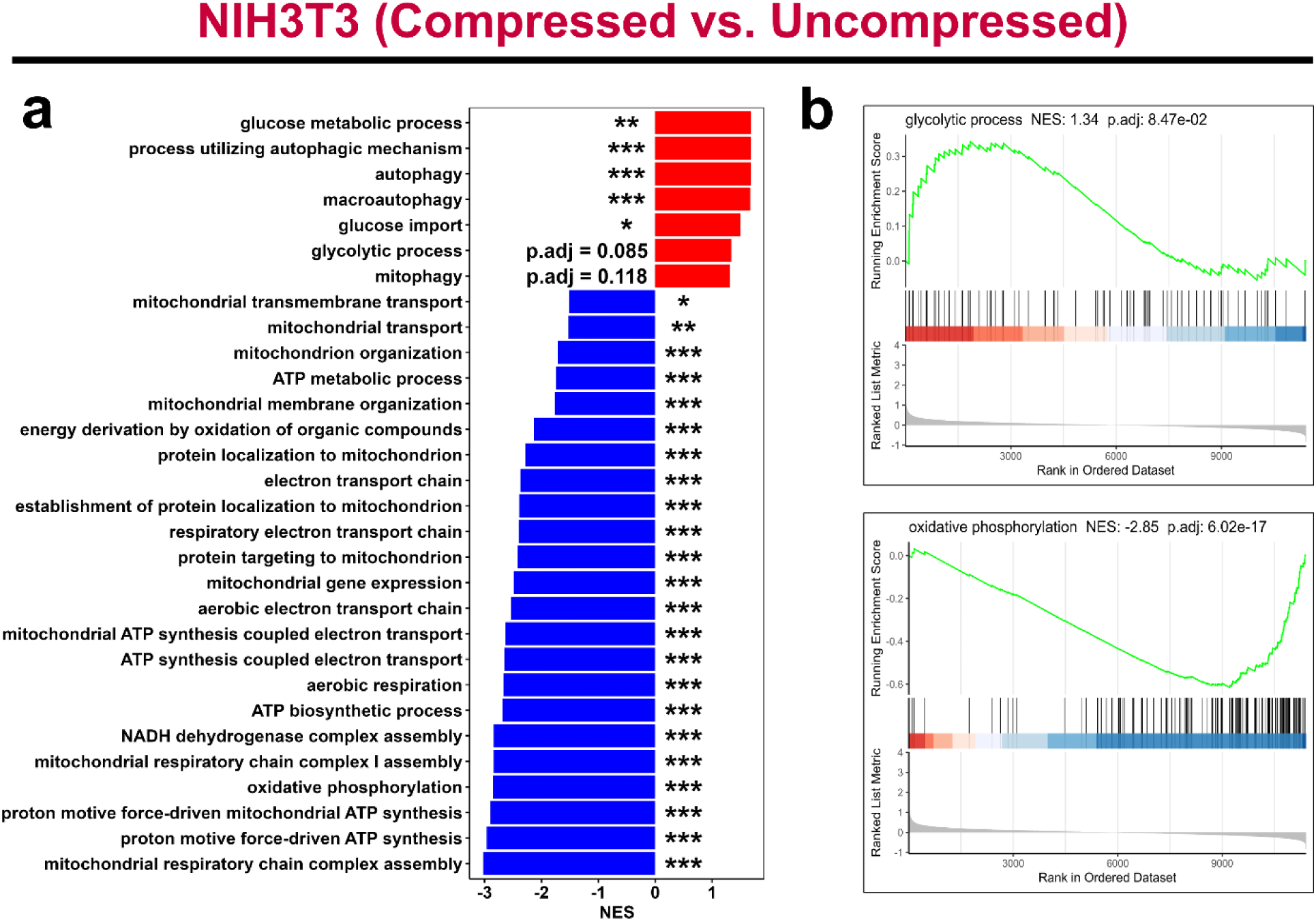
Compression alters the fibroblast transcriptome by upregulating key metabolic pathways that promote glycolysis while downregulating oxidative phosphorylation and related processes. Following 24 hours of *in vitro* compression, NIH3T3 fibroblasts were collected for transcriptomic analysis using bulk RNA-sequencing. **a)** Gene set enrichment analysis (GSEA) based on gene ontology (GO) terms shows that compressed fibroblasts (n = 3) downregulate oxidative phosphorylation and several related mitochondrial processes, as well as upregulate glycolysis and autophagic processes compared to their uncompressed counterparts (n = 3), confirming our FLIM results in Fig. 2. This highlights the sensitivity of instant FLIM as it can detect subtle metabolic changes that are reflected at the transcriptional level. **b)** GSEA plots from GO terms showing upregulation of glycolysis (top) and downregulation of oxidative phosphorylation (bottom) in compressed fibroblasts. Statistical significance was determined using permutation testing with Benjamini-Hochberg correction for multiple comparisons (\**p.adj* < 0.05, \*\**p.adj* < 0.01, \*\*\**p.adj* < 0.001). NES: normalized enrichment score.

### Compression induces increased mitochondrial fragmentation in undifferentiated adipocytes

We hypothesized that the compression-induced metabolic shifts in stromal cells would be accompanied by changes in mitochondrial morphodynamics. To investigate this, we stained the mitochondria of NIH3T3 fibroblasts and 3T3-L1 undifferentiated adipocytes with MitoTracker Red CMXRos following compression (**Fig. 5a,b**). Qualitative analysis of mitochondrial network morphology indicates a greater proportion of fragmented networks and swollen mitochondrion in compressed undifferentiated adipocytes (**Fig. 5b**), both of which are physical signs of mitochondrial dysfunction and damage.^31,32^ On the other hand, the mitochondria of compressed and uncompressed fibroblasts appear to be structurally similar (**Fig. 5a**).

**Figure 5.**
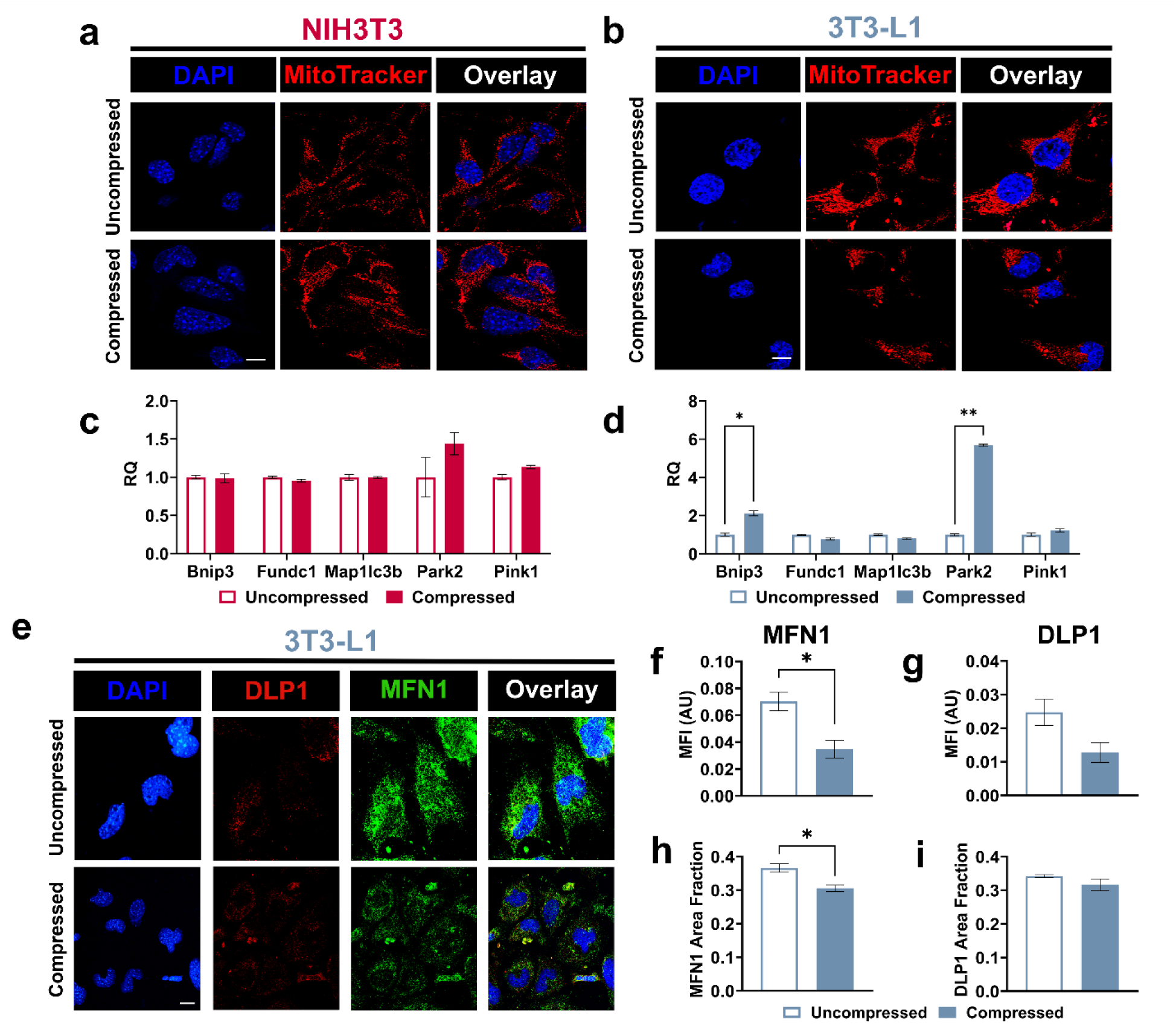
Compressed undifferentiated adipocytes exhibit altered mitochondrial morphodynamics with impaired fusion. Representative images of **a)** NIH3T3 fibroblasts and **b)** 3T3-L1 undifferentiated adipocytes stained with MitoTracker Red CMXRos. Qualitative observations of mitochondrial structure suggest that compression-induced mitochondrial damage occurs in undifferentiated adipocytes, but not fibroblasts, as indicated by greater fragmentation in the former. qPCR analysis of genes encoding proteins involved in mitophagy shows that none are differentially expressed in **c)** fibroblasts (n = 3), while *Bnip3* and *Park2* are significantly upregulated in **d)** compressed undifferentiated adipocytes (n = 3), confirming the qualitative data from **a)** and **b)**. **e)** Representative immunofluorescent images of 3T3-L1 cells stained for MFN1 (fusion marker) and DLP1 (fission marker). Immunofluorescent analysis reveals that the **f)** expression and **g)** area fraction (proportion of total cell area occupied by protein signal) of MFN1 is significantly lower in undifferentiated adipocytes subjected to compression (n = 3) compared to those that were not compressed (n = 3). On the other hand, there are no significant changes in DLP1 **h)** expression and **i)** area fraction between compressed (n = 3) and uncompressed (n = 3) 3T3-L1 cells. This suggests that the increased presence of mitochondrial fragments in compressed undifferentiated adipocytes likely results from impaired fusion dynamics. Error bars represent SEM and asterisks indicate statistical significance (\**p* < 0.05, \*\**p* < 0.01, \*\*\**p* < 0.001, \*\*\*\**p* < 0.0001) determined using a Student’s t-test. Scale bar is 10 µm.

### Mitophagy is upregulated in undifferentiated adipocytes in response to compressive stress

The selective degradation of dysfunctional/damaged mitochondria via mitophagy—a specialized form of autophagy—is crucial to the maintenance of mitochondrial homeostasis and, consequently, cell metabolism.^33^ Based on our observations of mitochondrial structure, we hypothesized that mitophagy is increased in compressed undifferentiated adipocytes but not in fibroblasts. Through qPCR analysis of mitophagy-related transcripts (*Bnip3*, *Fundc1*, *Map1lc3b*, *Pink1*, *Park2*), we found that *Bnip3* and *Park2* expression is upregulated in compressed undifferentiated adipocytes (**Fig. 5d**), signifying an activation of mitophagy mechanisms, while compressed NIH3T3 cells showed no significant changes in gene expression (**Fig. 5c**). The latter is consistent with our bulk-RNA sequencing analysis which revealed significant upregulation of general autophagy in compressed fibroblasts, but not mitophagy specifically (**Fig. 4a**).

### Undifferentiated adipocytes exhibit impaired mitochondrial fusion under compression

Given that mitochondrial network structure is carefully regulated by fission/fusion dynamics,^34^ we also evaluated the expression of DLP1 (fission protein) and MFN1 (fusion protein) in 3T3-L1 cells via immunofluorescence (**Fig. 5e**). By analyzing mean fluorescent intensity (MFI), we found that MFN1 (**Fig. 5f**) and DLP1 (**Fig. 5g**) expression is reduced in compressed cells, although this decrease is only significant for MFN1. Aligning with our MFI data, we also demonstrated that the area fraction (of total cell area occupied by protein signal) of MFN1 is significantly lower in compressed cells (**Fig. 5h**), whereas the area fraction of DLP1 remains relatively unchanged (**Fig. 5i**). These results suggest that breast tumor-mimicking *in vitro* compression impairs mitochondrial fusion, but not fission, dynamics in 3T3-L1 undifferentiated adipocytes. This likely explains the increased number of fragmented mitochondria in compressed cells, as a reduction in fusion events results in an imbalance in fission/fusion dynamics.

### Cancer cell-conditioned media has variable effects on compression-induced metabolic reprogramming of stromal cells

Stromal cells in the breast peritumor are also influenced by biological cues originating from nearby tumor cells. To incorporate these biochemical interactions and investigate the effect of mechanical vs. extreme (i.e., not physiological) biochemical cues on metabolic reprogramming, we compressed stromal cells in 100% 4T1 CM (i.e., media conditioned by murine 4T1 TNBC cells) and analyzed their metabolic state using instant FLIM (**Extended Data** Fig. 1a,b**,c**). Interestingly, our analysis revealed that adding 4T1 CM abrogates the effects of compression-induced metabolic reprogramming in NIH3T3 fibroblasts (**Extended Data** Fig. 1d,g) and 3T3-L1 undifferentiated adipocytes (**Extended Data** Fig. 1e,h), but not d3T3-L1 differentiated adipocytes (**Extended Data** Fig. 1f,i). Furthermore, we observed that cancer-conditioned media significantly increases lifetime in all cell types relative to those cultured in complete media regardless of compression status (**Fig. S4**). These results raise the question of whether biochemical versus compression-mediated metabolic changes are spatially dependent, with biochemical signals potentially dominating in some stromal cell types that may be located closer to the tumor interface, which we investigated next in patient samples.

### Metabolic reprogramming of peritumoral stromal cells is observed in human breast cancer that mimics in vitro compression results

To determine if stromal cell metabolism is spatially dependent, and to confirm our *in vitro* findings, we used instant FLIM to assess the metabolic state of adipocytes and fibroblasts in human TNBC and normal adjacent breast tissues. To identify and map the relative location of cell types of interest (i.e., fibroblasts and adipocytes), we stained serial tissue sections with H&E (**Fig. 6a**) and for α-SMA (**Fig. 6b**). Our results demonstrate that the metabolic activity of adipocytes, identified by their characteristic cell morphology, varies with their proximity to the tumor. Specifically, adipocytes within the tumor exhibit the shortest lifetime, while those in the normal stroma display the longest lifetime (**Fig. 6c,d**), indicating a shift towards glycolysis in tumor-proximal adipocytes. Fibroblasts exhibited similar spatial dependence of shorter (in tumor) to longer (in peritumor/normal) lifetimes. Interestingly, fibroblasts away from the tumor exhibit heterogeneous lifetimes likely owing to autofluorescence from overlapping and dense-ECM regions (**Fig. 6c,d**).

**Figure 6.**
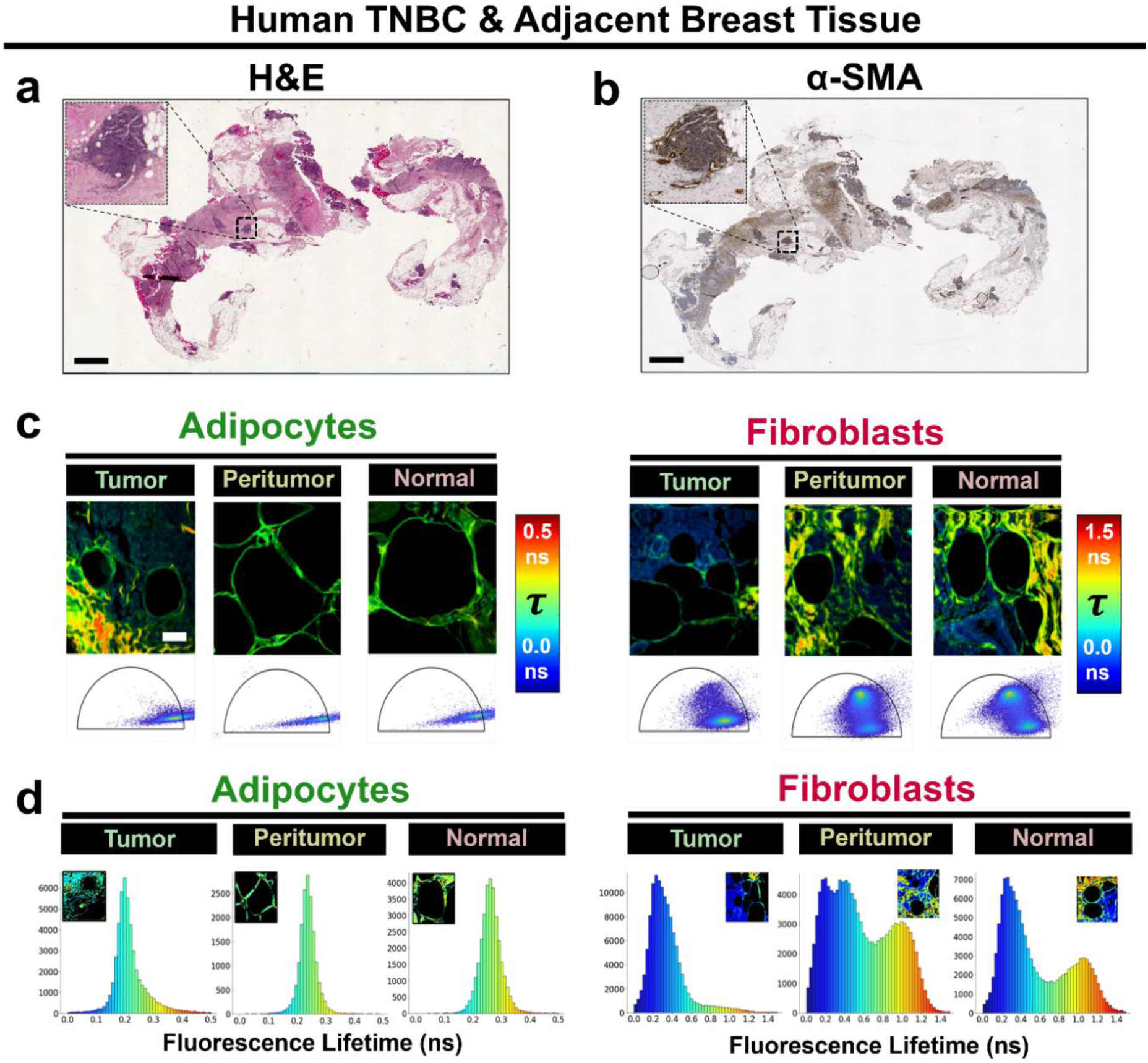
Fluorescence lifetime of human breast cancer stromal cells varies based on cell type and location relative to the tumor. Brightfield images of human TNBC with adjacent breast tissue stained **a)** with hematoxylin & eosin (H&E) and **b)** for α-SMA (fibroblast marker). **c)** Representative fluorescence lifetime images and associated phasor plot for mature adipocytes and fibroblasts in the tumor (left), peritumor (middle), and normal stroma (right). **d)** Distribution of fluorescence lifetime in tumor-associated (left), peritumoral (middle), and normal stromal (right) adipocytes and fibroblasts. Consistent with trends observed in Fig. 2f, histogram comparisons reveal that adipocytes in the tumor exhibit lower fluorescence lifetime values, indicative of a more glycolytic phenotype, whereas those in the normal stroma display higher values, suggesting greater oxidative metabolism. Adipocyte metabolism in the peritumor, on the other hand, appears intermediate between both regions, with cells likely undergoing a glycolytic transition potentially reflecting changes in maturity states. In fibroblasts, fluorescence lifetime trends in the peritumor and normal stroma are less distinct due to tissue heterogeneity, wherein multiple stromal components in the tissue contribute to a complex, non-unimodal lifetime distribution, but generally feature glycolytic trends. Advanced deconvolution methods are therefore needed to accurately resolve metabolic activity in fibroblasts from other tissue components (e.g., matrix) within the same region. Black scale bar represents 2 mm, white scale bar is 50 µm.

Furthermore, trends consistent with our *in vitro* and human *ex vivo* FLIM findings were evident in our analysis of DCN^+^ and PDGFRA^+^ fibroblasts and ADIPOQ^+^ and CIDEA^+^ adipocytes from human scRNA-seq and snRNA-seq data of breast cancer patient samples (**Fig. 7a**). To accomplish this, we determined the metabolic index (MI)—an in-house metric for characterizing the metabolic state of cells relative to the whole cell population—across different tissue zones (normal, peritumor, and tumor; **Fig. 7b**) for both cell types (**Fig. 7c**). Based on the definition of MI established here (see *Materials and Methods*), a higher value represents cells which are more glycolytic/less oxidative. We found that peritumoral fibroblasts and adipocytes have a higher mean MI than those originating from normal, healthy tissue; moreover, tumor-residing fibroblasts had the lowest mean MI (**Fig. 7d**). Thus, these results confirm that stromal cells harbor distinct metabolic profiles that are spatially dependent on tumor proximity, with peritumoral fibroblasts and adipocytes appearing to be more glycolytically active than their normal counterparts, largely mimicking our *in vitro* compression results. However, it remains to be determined to what extent this spatial metabolic reprogramming is a direct consequence of external tumor compression applied to the peritumor versus biochemical tumor signaling.

**Figure 7.**
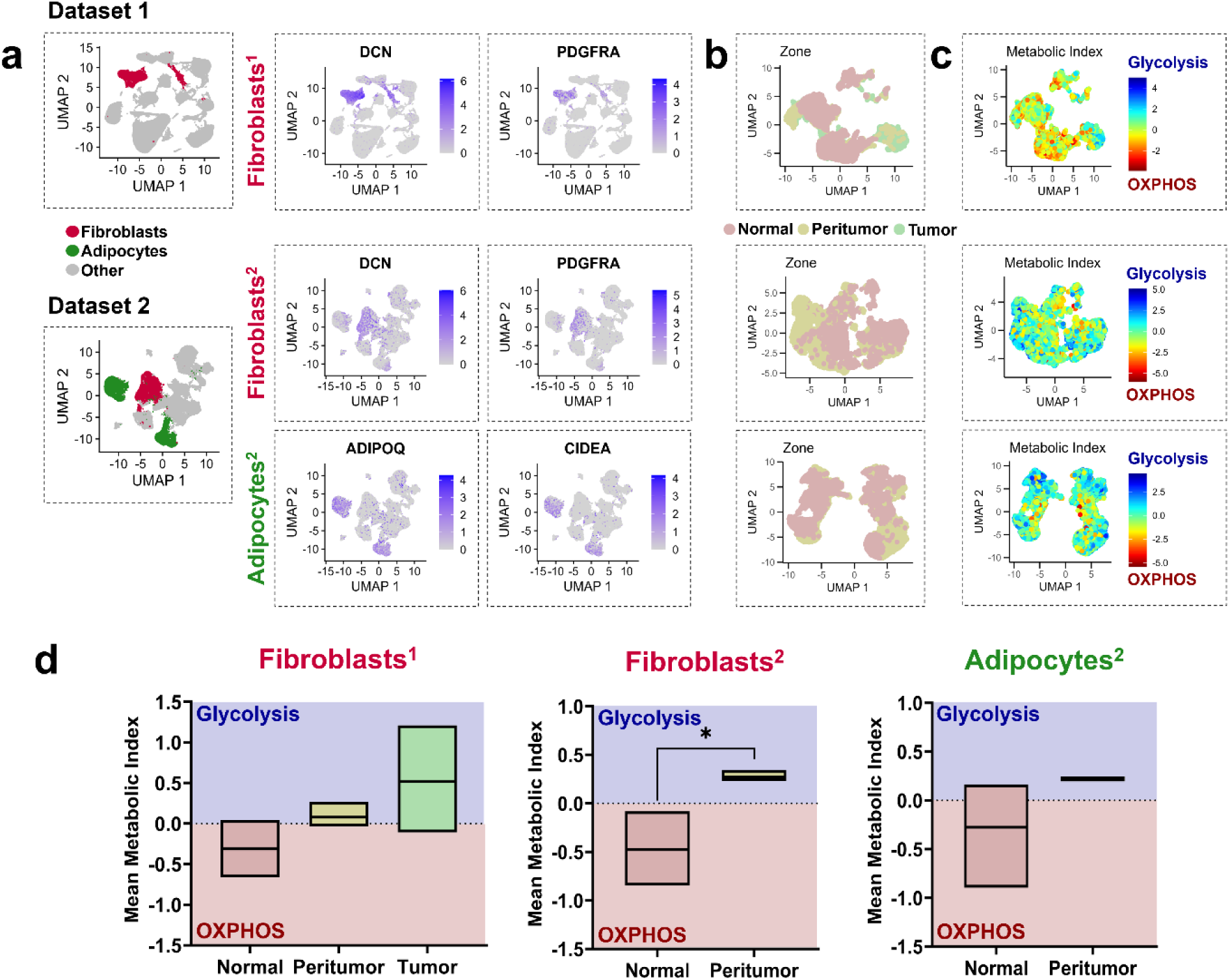
Human breast peritumoral fibroblasts and adipocytes are more glycolytic than normal stromal cells. To metabolically characterize human breast peritumoral adipocytes and/or fibroblasts, we conducted a retrospective analysis of two single-cell RNA sequencing datasets derived from human breast cancer patients. **a)** UMAP representation of fibroblasts (*DCN*^+^ and *PDGFRA*^+^) and/or adipocytes (*CIDEA*^+^ and *ADIPOQ*^+^) from dataset 1 (PRJNA960678; top) and dataset 2 (PRJNA932038; bottom) labeled by cluster. UMAP visualization of subclustered fibroblasts and/or adipocytes labeled according to their **b)** spatial location within the tissue (normal, peritumor, or tumor tissue zones) and **c)** metabolic index (MI). **d)** Mean MI analysis per patient reveals that stromal cells in the peritumor region exhibit higher glycolytic activity than their normal counterparts, with MI increasing as proximity to the tumor increases. The MI is a relative measure of metabolic activity based on the normalization and standardization of raw metabolic scores. Here, MI > 0 indicates cells are more glycolytic while MI < 0 suggests they have higher levels of oxidative phosphorylation activity. Statistical significance for dataset 1 was determined using a Student’s t-test while significance for dataset 2 was determined using one-way ANOVA with Tukey’s HSD correction. Asterisks indicate significance levels (\**p* < 0.05, \*\**p* < 0.01, \*\*\**p* < 0.001, \*\*\*\**p* < 0.0001). ^1^PRJNA960678; ^2^PRJNA932038.

## Discussion

The use of gene expression-based molecular profiling has redefined the field’s view of the breast peritumor. Notably, studies have shown that the peritumor possesses a unique molecular signature that distinguishes it from healthy mammary and malignant tumor tissue, despite often appearing morphologically normal.^6-9,35^ These distinct characteristics reflect active remodeling of the peritumor driven by dynamic tumor-host interactions. Importantly, this tumor-induced remodeling alters the functional landscape of the surrounding mammary tissue, transforming it into a microenvironment that supports tumor growth and progression.^4,36-39^ Thus, the peritumor serves as a critical hub of biomarkers which can provide valuable insight into key clinical metrics, including early prognosis, relapse prediction, and therapeutic response, as well as an attractive target for novel therapeutic development.^6,9,35,40^

While these molecular signatures have underscored the distinctiveness of the peritumor, the mechanisms underlying the acquisition of these traits remains limited. Recent work has implicated the tumor in disrupting the surrounding tissue at the cell and tissue length scales through both biochemical (e.g., chemokine signaling) and biophysical/mechanical (e.g., solid stresses) means.^3,19,39^ Such disruption can lead to physical changes—including altered stiffness, morphology, and microarchitecture, as well as compression of the stroma—in the breast peritumor that induce and reinforce tumorigenic cell phenotypes via metabolic reprogramming. Indeed, differences in matrix stiffness and collagen density have been observed to directly regulate key attributes of malignancy (e.g., migration, invasion, and proliferation) in breast epithelial and cancer cells cultured in 2D and 3D environments by tuning their bioenergetics.^41-47^ Mammary stromal cell metabolism is also altered in response to these mechanical aberrancies, wherein stiffer matrices appear to increase glycolytic flux and soluble factor production in normal fibroblasts resulting in enhanced breast epithelial cell migration.^16^

In contrast, while much is known about how peritumoral matrix properties may influence cell metabolism, significantly fewer studies have focused on the metabolic response of the breast pTME to solid stress. A physical hallmark of cancer, solid stress arises in part from the reciprocal mechanical interactions between the growing tumor and the surrounding host tissue, which resists deformation caused by growth-induced tumor compressive forces.^18,19^ Initial studies looking at the effects of solid stress *in vitro* have demonstrated that it can enhance the migration and invasiveness of breast cancer cells.^27^ Furthermore, reducing solid stress in TNBC mouse models has been shown to potentiate chemotherapy^48^ and immunotherapy.^49^ The effects of solid stress on stromal cells have also been studied, with previous work demonstrating that compression drives a myriad of phenotypic and functional changes that support tumor growth. These changes include the de-differentiation of adipocytes into multipotent stem cells,^39^ activation of normal human pancreatic fibroblasts,^29^ spreading and enwrapment of cancer cells by murine fibroblasts,^50^ and increased microRNA expression^51^ and metabolic changes in human breast CAFs.^52^ Despite these insights, the impact of tumor-generated compressive forces on the metabolism of normal host tissue cells remains largely unexplored.

To address this gap, we investigated the metabolic response of stromal cells—namely, fibroblasts and adipocytes—found in the breast peritumor when subjected to breast tumor-mimicking compression by examining optical, transcriptional, and functional readouts of metabolic state. Importantly, while fibroblasts are well known for their mechanosensitivity, relatively few studies have explored the mechanoresponsive properties of adipocytes,^53-55^ which are highly relevant in the context of breast cancer. Furthermore, it should be noted that while undifferentiated adipocytes are relatively scarce in the normal mammary stroma except during pregnancy and lactation;^56,57^ the onset of breast tumorigenesis increases the abundance of undifferentiated-like adipocytic cells by promoting the de-differentiation of resident adipocytes in the tumor-adjacent host tissue.^58,59^ These de-differentiated cells can further transform into cancer-associated adipocytes by undergoing transcriptional changes and metabolic reprogramming, fueling tumor growth, progression, and metastasis.^58-60^

The cornerstone of our study was fluorescence lifetime imaging microscopy (FLIM), a cutting-edge tool that has been previously used for *in vitro*, *ex vivo*, and *in vivo* metabolic characterization of biological materials, including breast cancer spheroids,^45^ stromal macrophages,^61^ malignant and benign tumors,^62,63^ and mono- and co-cultures of cancer cells and/or fibroblasts.^64^ Notably, we specifically utilized our recently developed and novel FD-FLIM technique, instant FLIM, due to its improvements in image acquisition and data processing speed, SNR, cost-efficiency, and ease of implementation—common limitations encountered in TD-FLIM and/or FD-FLIM systems.^20,23^ In accordance with the aforementioned studies, we demonstrated that compression-induced metabolic changes are cell-type specific and environmentally dependent. Notably, compressed fibroblasts and undifferentiated adipocytes exhibited metabolic shifts toward a more glycolytic or oxidative state, respectively, when cultured in DMEM, but not in cancer-conditioned media. In contrast, differentiated adipocytes seemingly underwent glycolytic reprogramming in response to compression regardless of culture conditions. It is important to note that these metabolic classifications may be an oversimplification. Our interpretations were based on changes in the fluorescence lifetime of the metabolic cofactor, NADH. As discussed by Szulczewski et al., the spectra of NADH and its functionally promiscuous anabolic counterpart, NAD(P)H,^65^ are similar, making it difficult to accurately resolve their relative contributions.^61^

Nevertheless, we utilized instant FLIM because it offers the unique advantage of being label-free, non-destructive, and spatially resolved, thus allowing users to detect subtle, yet significant, molecular changes *in situ*. These advantages are exemplified here, with our bulk-transcriptomic analysis of compressed fibroblasts revealing metabolic gene expression changes consistent with our FLIM data; this correlation emphasizes the sensitivity of instant FLIM to transcriptional changes. Additionally, the system is broadly compatible as it can accommodate a variety of sample types. Alongside our *in vitro* studies, we were able to demonstrate using paraffin-embedded human breast cancer tissues that stromal cell metabolism is spatially dependent, with cells becoming increasingly glycolytic as their proximity to the tumor increases. It is worth noting that, due to the heterogeneous landscape of the peritumor and normal stroma, we were unable to distinguish the lifetime contribution of single-cell fibroblasts from other cellular and non-cellular tissue components (**Fig. 7c,d**). Therefore, future experiments implementing advanced deconvolution methods will be necessary to accurately resolve and characterize fibroblast metabolic state.

Importantly, although we did not observe differences in metabolic activity through our analysis of real-time ATP production, we attribute this to experimental and sensitivity limitations. In particular, our system utilizes transwell chambers to ensure there is adequate gas and nutrient exchange during compression. However, after compression, cells must be enzymatically detached, replated in a Seahorse XF Analyzer-compatible plate, and incubated for at least one hour before real-time metabolic measurements can be collected. These time-consuming and stressful pre-processing steps introduce external variables that potentially alter cellular metabolism in a way that is no longer representative of the original metabolic state. Nevertheless, our results collectively show that applied compressive stress promotes distinct metabolic changes. These findings also demonstrate the advantage of using instant FLIM over traditional metabolic profiling methods, as it delivers superior sensitivity while requiring minimal sample processing.

This study is, to the best of our knowledge, the first of its kind to directly evaluate the mechano-metabolic response of non-malignant stromal cells commonly present in the mammary host tissue. While our investigation was limited to fibroblasts and adipocytes, similar studies are underway for other stromal cell types, including immune and vascular cells. Another limitation in our study is that our conditioned media experiments did not faithfully recapitulate *in vivo* conditions. In addition to breast pTME stromal cells, both cancer cells and stromal cells within the tumor microenvironment (TME) experience compressive stress, which can promote the secretion of additional biochemical factors into the pTME. Therefore, solid stress may also indirectly drive tumor-host mechano-metabolic crosstalk by first reprogramming the TME.

Furthermore, several questions remain unanswered, including the mechanisms governing the differential responses to compression that we observed. One possibility may lie in our finding that both compressed fibroblasts and undifferentiated adipocytes exhibited YAP activation, a mechanotransducer whose metabolic effects are known to vary. For instance, YAP promotes glycolysis in cardiomyocytes in response to acute pressure overload^66^ and enhances oxidative phosphorylation in regulatory T cells cultured on stiff matrices.^67^ Moreover, our results revealed that compression enhances mitochondrial fragmentation, increases the expression of the mitophagy markers *Park2* and *Bnip3*, and reduces the level of the mitochondrial fusion protein MFN1 in undifferentiated adipocytes. The observed increase in mitophagy may be a consequence of elevated oxidative phosphorylation, which can generate reactive oxygen species and contribute to mitochondrial oxidative stress.^68,69^ Furthermore, the reduction in mitochondrial fusion may be linked to increased parkin (*Park2*) expression, an E3 ubiquitin ligase that targets MFN1 for degradation.^70^ However, further studies are required to elucidate these mechanistic relationships. Other areas of interest for future investigation include regulation by mechanosensitive signaling^71^ and changes in the actin cytoskeleton.^72^ Additionally, *in vivo* causal studies investigating mammary tissue response to compression (e.g., with a compressive window^73,74^) would further confirm our observations and establish a correlation with our human data.

Taken together, these findings offer new insights into tumor-host crosstalk, demonstrating that tumor-induced compressive forces play a crucial role in shaping the metabolism of host tissue cells. This phenomenon may extend to other solid malignancies and disease contexts where solid masses form (e.g., granulomas^75,76^ or benign tumors^77^). Importantly, this biophysical crosstalk presents an area of untapped translational potential. Consequently, future studies focused on elucidating mechano-metabolic abnormalities in the peritumoral microenvironment of the breast could be pivotal in identifying novel, targetable vulnerabilities to enhance therapeutic response.

## Methods

### Cell Culture

Murine NIH3T3 fibroblasts (American Type Culture Collection [ATCC], CRL-1568, Manassas, VA, USA) and 3T3-L1 undifferentiated adipocytes (ATCC, CRL-173) were grown in the following complete media: Dulbecco’s Modified Eagle Medium (DMEM; Corning, 10-03-CV, Corning, NY, USA) supplemented with 10% fetal bovine serum (FBS; Corning, 35-015-CV) and 1X penicillin-streptomycin (P/S; Corning, 30-002-CI). Cells were maintained in a 37 °C, 5% CO_2_ humidified incubator and passaged once they reached ∼70% confluency.

### Adipocyte Differentiation

Low passage (< P6) 3T3-L1 undifferentiated adipocytes were seeded into clear 0.4 μm 6-well transwell inserts (Sterlitech, 9300412, Auburn, WA, USA) in a 6-well plate at a density of 9.3 x 10^3^ cells/cm^2^, grown in complete media, and maintained in a 37 °C, 5% CO_2_ humidified incubator. Once the cells reached ∼80-90% confluency, they were differentiated based on a previously described protocol.^78^ Briefly, cells were treated for 48 hours with a mixture of 1 μg/ml insulin (Sigma-Aldrich, 10516, St. Louis, MO, USA), 0.25 μM dexamethasone (Sigma Aldrich, D4902), 0.5 mM 3-isobutyl-1-methylxanthine (Sigma-Aldrich, I5879), and 2 μM rosiglitazone (Sigma-Aldrich, R2408), followed by 10 consecutive days of treatment with 1 μg/ml insulin. After 10 days, the differentiated 3T3-L1 (d3T3-L1) adipocytes were ready to be compressed.

### Generation of Conditioned Media

Murine 4T1 triple-negative breast cancer (TNBC) cells (ATCC, CRL-2539) were cultured in Roswell Park Memorial Institute (RPMI) medium (Corning, 10-040-CV) supplemented with 10% FBS and 1X P/S. Once cells reached ∼80% confluence, they were washed once with 1X PBS to remove traces of serum and fresh serum-free RPMI was added. After 48 hours, the 4T1 conditioned media (CM) was collected, centrifuged for 10 minutes at 1000 xg, sterile filtered using 0.22 μm syringe filters (NEST Scientific USA, 380121, Woodbridge, NJ, USA), separated into aliquots, and stored in -80 °C until they were ready to be used.

### In Vitro Transwell Compression Assay

To study the effects of solid stress *in vitro*, we utilized a previously established transwell compression assay (**Fig. 1a**).^27-29^ Briefly, ∼2 x 10^5^ NIH3T3 or 3T3-L1 cells were seeded in clear 0.4 μm 6-well transwell inserts and allowed to attach overnight in complete media. The 0.4 μm pore size was chosen to impede cell migration while also permitting the transport of oxygen and nutrients to the cells.^29^ The following day, the media was replaced with fresh complete media or 4T1 CM. Following media replacement, a 1% (w/v) agarose cushion was carefully placed on top of the cells (NIH3T3, 3T3-L1, or d3T3-L1) to prevent contact between the weight and cells. Agarose cushions were prepared using serum-free DMEM and equilibrated the previous night in serum-free DMEM in a 37 °C, 5% CO_2_ humidified incubator. Finally, a biocompatible, 3D-printed polylactic acid weight was placed on top of the cushion to apply ∼0.14 kPa of compressive stress—the approximate magnitude of solid stress exerted by murine breast tumors^79^—to the cells for 24 hours. Wells without added weight served as control wells.

### Sample Preparation for Imaging

After 24 hours of compression, all weights and agarose cushions were carefully removed from the wells. Cells were then washed 3 times with 1X PBS, fixed in 4% paraformaldehyde (Thermo Fisher Scientific, J19943.K2, Waltham, MA, USA) for 15 minutes, and washed 3 more times with 1X PBS. Following this, the unstained cells were either immediately mounted onto SuperFrost microscope slides (VWR, 48311-703, Radnor, PA, USA) for fluorescence lifetime imaging, or stained then mounted. To properly mount the cells, the membrane of the transwell insert was removed, cut at the edges into a rectangle to avoid wrinkling, and loaded flat onto the microscope slide.

### Human Triple-Negative Breast Cancer Tissues

Human triple-negative breast cancer tissue samples were obtained from the Harper Cancer Research Institute’s (South Bend IN, USA) Biosample Repository with Institutional Review Board approval (Protocol #21-01-6380). To visualize general cellular structures, tissues were stained with hematoxylin and eosin (H&E). Immunohistochemistry for α-smooth muscle actin (α-SMA) expression was also performed to identify fibroblasts. Whole slide brightfield images were acquired using the Keyence BZ-X810 (Keyence, Itasca, IL, USA).

### Fluorescence Lifetime Imaging

Here, we use “instant FLIM," a custom-built two-photon excited frequency-domain FLIM (FD-FLIM) which incorporates a mode-locked femtosecond-pulsing Ti:Sapphire laser (Spectra-Physics Mai Tai BB; 710–990 nm, 100 fs, 80 MHz; **Fig. 1b**).^23^ For all samples (NIH3T3, 3T3-L1, d3T3-L1, and human breast cancer tissue), 800 nm two-photon excitation was selected and a dynamic field of view was applied. Fluorescence lifetime analysis was performed using the phasor approach, a fit-free method for segmentation based on the phasor components g (cosine) and s (sine) of the fluorescence signal in the phasor plot.^80-82^ For *in vitro* samples, lifetime values are averaged by applying a mask from the intensity map to g and s for averaged g and s due to sample consistency across different FOVs. In human breast cancer tissue, a pure-lifetime map with a corresponding histogram under a same color bar is used to differentiate fibroblasts (α-SMA stained) from adipocytes.

### Cell Staining and Analysis

Fixed cells were permeabilized with 0.1% Triton X-100 (Thermo Fisher Scientific, 0694) for 10 minutes, blocked with 2% bovine serum albumin (VWR, 0332) or 10% normal goat serum (Sigma-Aldrich, G6767) for 60 minutes, then incubated with the following primary antibodies overnight at 4 °C: mouse anti-DLP1 (1:300; BD Biosciences, 611112, Franklin Lakes, NJ, USA) and rabbit anti-MFN1 (1:500; Proteintech, 13798-1-AP, Rosemont, IL), and mouse anti-YAP Alexa Fluor 488 (1:100; Santa Cruz Biotechnology, sc-376830, Dallas, TX, USA). The following day, cells were incubated with goat anti-rabbit 488 (1:500; Abcam, ab150077, Cambridge, UK), goat anti-mouse 594 (1:500; Thermo Fisher Scientific, A-11005), and/or counterstained with DAPI (1:1000, Thermo Fisher Scientific, D1306) for 45 minutes at room temperature. The samples were then mounted onto microscope slides and imaged using a Nikon AX-R confocal microscope (Nikon, Minato City, Tokyo, JPN). YAP localization was characterized using CellProfiler by determining the logarithmic ratio of YAP intensity in the nucleus to that in the perinuclear space (log[nuclear:perinuclear intensity]). The expression and area fraction of DLP1 and MFN1 were analyzed using ImageJ. Additionally, live cells were stained with 150 nM MitoTracker Red CMXRos (Thermo Fisher Scientific, M7512) for 20 minutes then fixed and stained with DAPI (1:1000) to assess mitochondrial network morphology. To obtain high-resolution images of the mitochondria, cells were imaged at 60x using the ZEISS LSM 900 with Airyscan 2 (ZEISS, Oberkochen, Baden-Württemberg, GER).

### Seahorse Metabolic Analysis

To evaluate relative ATP production via glycolysis and oxidative phosphorylation, a Real-Time ATP Rate Assay (Agilent Technologies, 103592-100, Santa Clara, CA) using the Seahorse XFe96 Extracellular Flux Analyzer (Agilent Technologies) was conducted according to the manufacturer’s protocol. Briefly, after 24 hours of compression, cells were carefully decompressed, enzymatically detached, centrifuged, and resuspended in Seahorse XF DMEM (Agilent Technologies, 103575-100) supplemented with glucose (10 mM; Agilent Technologies, 103577-100), pyruvate (1 mM; Agilent Technologies, 103578-100), and glutamine (2 mM; Agilent Technologies, 103579-100). Cells were then replated in PDL-coated Seahorse microplates (Agilent Technologies, 103798-100) at the following densities: 2.78 x 10^4^ cells/well for NIH3T3 fibroblasts and 2.07 x 10^4^ cells/well for 3T3-L1 undifferentiated adipocytes. Following this, the plates were centrifuged at 200 xg for 1 minute to allow cells to adhere to the bottom of the well and incubated for 3 hours at 37 °C in a non-CO_2_ incubator. The oxygen consumption rate (OCR) was measured then over time following sequential injections of 1.5 μM oligomycin and 0.5 μM rotenone + antimycin A (Rot/AA). For normalization, cells were fixed in 4% PFA for 15 minutes, stained with DAPI (1:1000), and fluorescent intensity of each well was measured using a Tecan Plate Reder (Tecan, Morrisville, NC, USA) immediately after the run. Analysis was performed using Agilent Seahorse Analytics.

### Glucose Uptake Study

The Glucose Uptake-Glo™ Assay (Promega, J1341, Madison, WI, USA) was used to determine whether differences in glucose uptake exist between compressed and uncompressed cells. The manufacturer’s recommended protocol for cells treated in 6-well plates was followed. Briefly, cells were treated with 2DG for 10 minutes, lysed with Stop Buffer, transferred to a 96 well black clear bottom plate (Greiner Bio-One, 655096, Frickenhausen, GER), neutralized with Neutralization Buffer, then treated with 2DG6P detection reagent. Importantly, instead of treating the cells *in situ* with 2DG solution following decompression, the cells were enzymatically detached, centrifuged, and resuspended in 2DG so that they could be counted. After 1 hour of incubation with the detection reagent, luminescence was read using a Tecan Plate Reader and normalized to the estimated cell count per well for each sample.

### qPCR

TaqMan quantitative PCR (qPCR) was conducted to evaluate the expression of transcripts associated with mitophagy and response to mechanical stress. To do this, cells were collected after 24 hours of compression via cell scraping, centrifuged, and lysed in 300 μl TRI Reagent (Zymo Research, R2052, Irvine, CA, USA). RNA purification was then carried out using Zymo’s Direct-zol^TM^ RNA Miniprep Kit (Zymo Research, R2052) followed by cDNA synthesis using the High-Capacity cDNA Reverse Transcription Kit (Thermo Fisher Scientific, 4374966). The relative gene expression of *Bnip3* (Mm01275600_g1), *Fundc1* (Mm00511132_m1), *Map1lc3b* (Mm00782868_sh), *Park2* (Mm01323528_m1), *Pink1* (Mm00550827_m1), *Ctnnb1* (Mm00483039_m1), *Piezo1* (Mm01241549_m1), *Trpv4* (Mm00499025_m1), and *Yap1* (Mm01143263_m1) were evaluated using the 2^−ΔΔCt^ method normalized to *Gapdh* (Mm99999915_g1) expression. All TaqMan probes were purchased from Thermo Fisher Scientific.

### Bulk RNA-sequencing Analysis

Following RNA purification as described above, RNA quality and integrity was determined using the Qubit Fluorometer and Agilent 2100 Bioanalyzer system; all samples had a RIN > 8.0. Samples that passed quality control were used to generate 150 bp paired-end libraries with the NEBNext Ultra II Directional RNA Library Prep Kit (New England Biolabs, E7760, Ipswich, MA, USA). Libraries were sequenced at the Indiana University Center for Medical Genomics on the NovaSeq 6000 using the S4 flow cell, with a target depth of 37 million read per sample. Reads were aligned using the STAR tool. Differential gene expression analysis was performed using DESeq2.^83^ Gene set enrichment analysis (GSEA) for gene ontology (GO) terms was conducted using gseGO and visualized using gseaplot2 from the clusterProfiler package.^84^

### Single-Cell RNA Sequencing Analysis

scRNA-seq and snRNA-seq data from human breast cancer patients was obtained from Sequence Read Archive (PRJNA960678^85^ and PRJNA932038^86^). Fibroblasts (PRJNA960678 and PRJNA932038) were subclustered based on the differential expression of *DCN* and *PDGFRA* while adipocytes (PRJNA932038) were sublcustered based on the differential expression of *CIDEA* and *ADIPOQ*. Raw metabolic scores for glycolysis (GLY) and oxidative phosphorylation (OXPHOS) were computed using the scMetabolism package.^87^ A power transformation was then applied to each set of scores to normalize the data. To assess the metabolic state of individual cells, a metabolic index (MI) was created by subjecting the transformed scores to robust z-score standardization and subtracting the resulting values from each other as shown below:

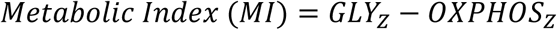

where GLY_Z_ and OXPHOS_Z_ are robust z-standardized glycolysis and oxidative phosphorylation scores, respectively. Therefore, relative to the whole cell population, MI < 0 indicates a shift towards higher oxidative phosphorylation while cells with MI > 0 are becoming more glycolytic.

### Statistical Analysis

Statistical analysis and data visualization was carried out using R and GraphPad Prism. In data reported for FLIM of cell samples, Seahorse, qPCR, glucose uptake assay studies, and immunofluorescence datasets errors bars, represent mean ± standard error of the mean (SEM) with significance being determined using a Student’s t-test. GSEA enrichment scores from bulk RNA-sequencing data were evaluated for statistical significance through permutation testing, with multiple comparisons corrected by the Benjamini-Hochberg method (default settings for gseGO). The significance of the mean metabolic index determined from scRNA-seq datasets was assessed using one-way ANOVA with Tukey’s HSD correction for PRJNA960678 and Student’s t-test for PRJNA932038. The level of significance is represented by the following convention unless otherwise specified: \**p* < 0.05, \*\**p* < 0.01, \*\*\**p* < 0.001, \*\*\*\**p* < 0.0001.

## Acknowledgements

We thank Dr. Yichun Wang and Ms. Gaeun Kim for their generous donation of NIH3T3 fibroblasts and Dr. Donny Hanjaya-Putra for use of his confocal microscope. We also thank Ms. Jensen Amens for providing guidance on adipocyte differentiation and Ms. Ellie Johandes for her insight on the glucose uptake assay. Human tissue samples were obtained with the help of Ms. Toni Mayberry and Dr. Laurie Littlepage through the Haper Cancer Research Institute Tissue Biorepository under protocols approved by the Institutional Review Board (IRB) of the University of Notre Dame (Protocol #21-01-6380). All procedures were conducted in accordance with institutional guidelines and the principles outlined in the Declaration of Helsinki. Histological staining of human breast cancer tissue samples was conducted in the Notre Dame Histology Core, and whole-slide scans of stained tissue samples were taken in the Notre Dame Integrated Imaging Facility. We thank Dr. Sarah Chapman and Dr. Sara Cole for their knowledge and expertise as well as time towards this research. We also thank Ms. R’nld Rumbach for her technical assistance. This work was supported by the National Institutes of Health (NIH/NIGMS R35GM151041 to M.D. and NIH/NCI R01CA275423 to P.Z.).

## Contributions

**Julian Najera:** conceptualization, data curation, formal analysis, investigation, methodology, software, validation, visualization, writing – original draft, writing – review and editing; **Hao Chen:** conceptualization, data curation, formal analysis, investigation, methodology, software, validation, visualization, writing – original draft, writing – review and editing; **Bianca Batista:** formal analysis, investigation, visualization, writing – review and editing; **Frank Ketchum:** investigation, validation, visualization, writing – review and editing; **Aktar Ali:** investigation, resources, supervision, validation, writing – review and editing; **Pinar Zorlutuna:** funding acquisition, resources, writing – review and editing; **Scott Howard:** conceptualization, methodology, project administration, resources, software, supervision, validation, writing – review and editing; **Meenal Datta:** conceptualization, funding acquisition, methodology, project administration, resources, software, supervision, validation, writing – review and editing.

## Data Availability

Data and codes are available upon request.

---

**Extended Data Figure 1.**
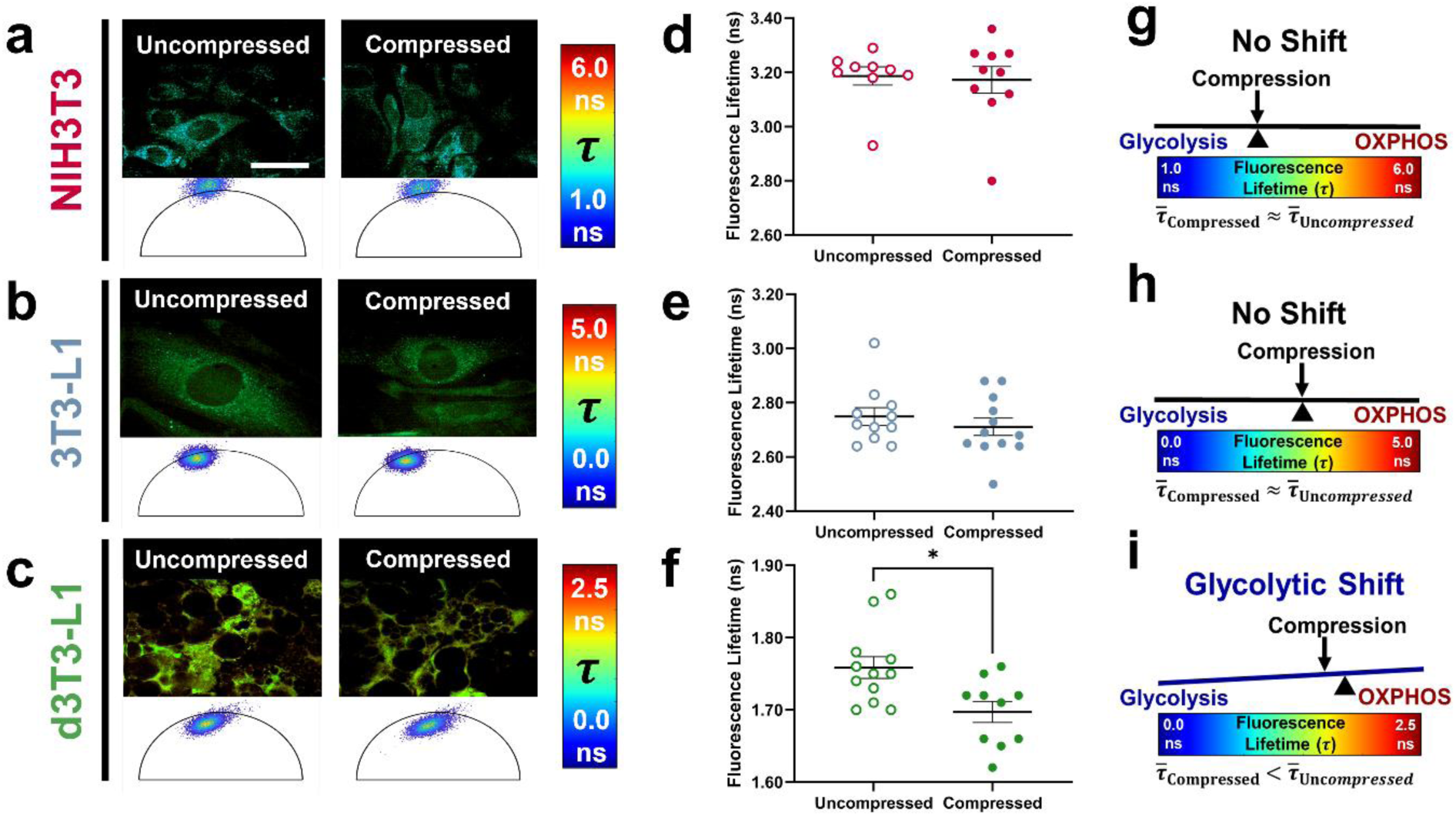
Biochemical effects from breast cancer conditioned media (CM) overcomes compression-induced metabolic reprogramming in fibroblasts and undifferentiated adipocytes, but not differentiated adipocytes. Representative fluorescence lifetime images (top) and phasor plots (bottom) of **a)** NIH3T3 fibroblasts (n = 9 uncompressed, 10 compressed), **b)** 3T3-L1 undifferentiated adipocytes (n = 11 uncompressed, 12 compressed), and **c)** d3T3-L1 adipocytes (n = 12 uncompressed, 10 compressed) cultured in murine 4T1 triple-negative breast cancer cell CM. Fluorescence lifetime analysis shows that the mean lifetime (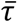) does not change in **d)** NIH3T3 fibroblasts and **e)** 3T3-L1 undifferentiated adipocytes following compression in 100% 4T1 CM. This indicates there is no shift in metabolic state in both cell types as shown in **g)** and **h)**. Conversely, fluorescence lifetime is significantly lower in **f)** compressed d3T3-L1 adipocytes suggesting a **i)** glycolytic shift as in Fig. 2f**,i**. Thus d3T3-L1 cells may be more metabolically sensitive to mechanical (rather than biochemical) signals. These results raise the possibility that, in some peritumoral stromal cell types, the metabolic response to growth-induced tumor compressive forces may not only be influenced by cell identity, but also spatial proximity to tumor where biochemical cues are more likely to dominate closer to the tumor. Error bars represent SEM and asterisks indicate statistical significance (\**p* < 0.05, \*\**p* < 0.01, \*\*\**p* < 0.001, \*\*\*\**p* < 0.0001) determined using a Student’s t-test. Scale bar is 50 µm.

**Figure S1.**
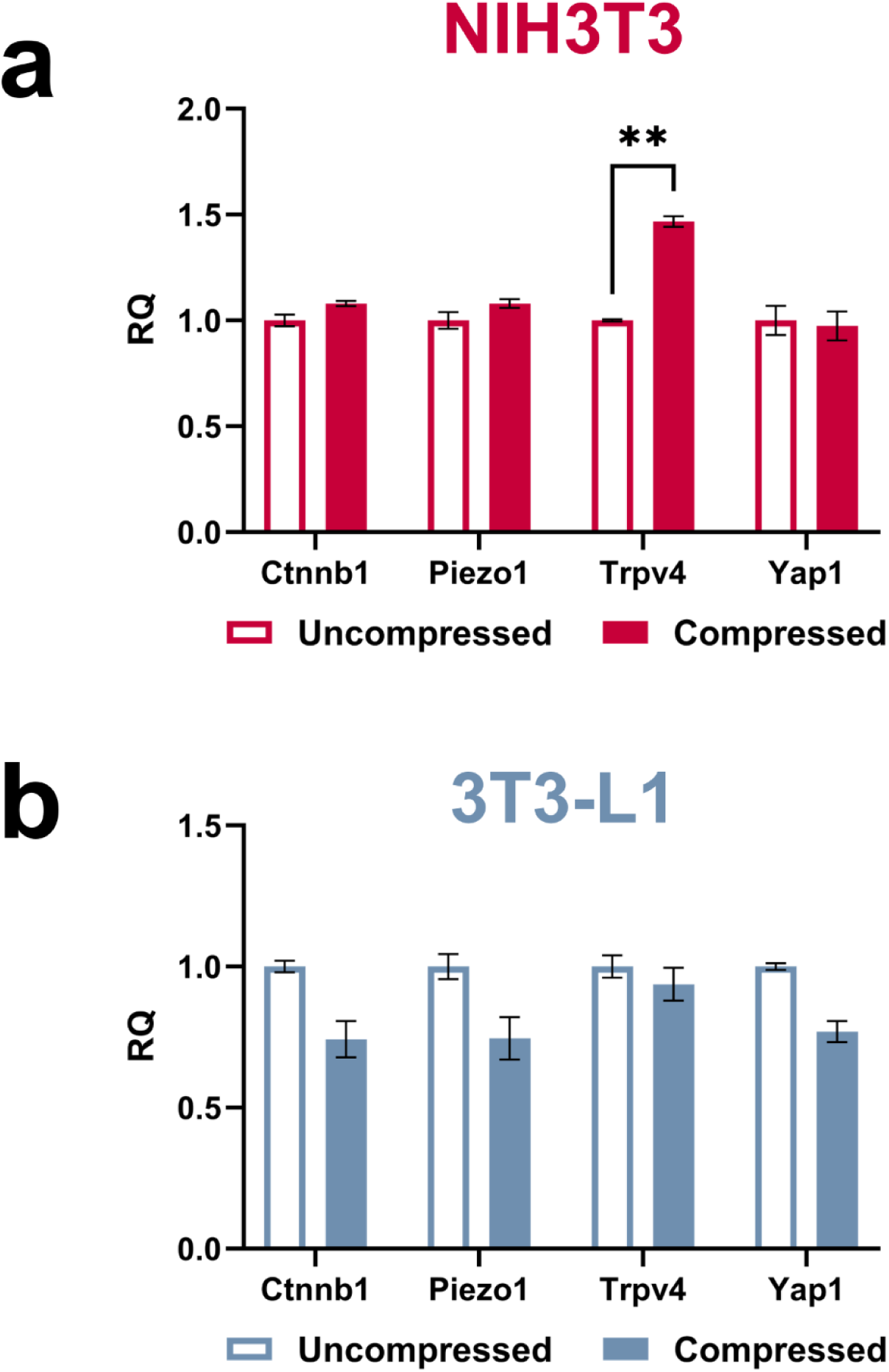
Compression does not widely alter the expression of mechanotransduction-related transcripts. Gene expression analysis of markers involved in mechanotransduction via qPCR shows that *Trpv4* expression is moderately upregulated in **a)** compressed NIH3T3 fibroblasts (n = 3), but not **b)** compressed 3T3-L1 undifferentiated adipocytes (n = 3). On the other hand, *Ctnnb1*, *Piezo1*, and *Yap1* expression in both cell types is not significantly affected by compressive stress. Error bars represent SEM and asterisks indicate statistical significance (\**p* < 0.05, \*\**p* < 0.01, \*\*\**p* < 0.001, \*\*\*\**p* < 0.0001) determined using a Student’s t-test. RQ: relative quantification.

**Figure S2.**
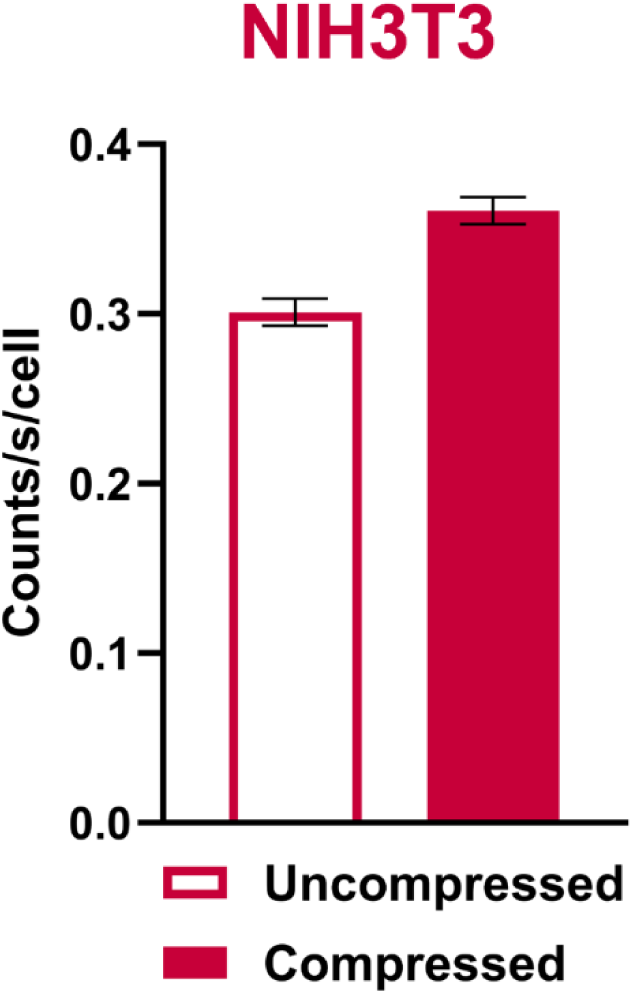
Compression enhances glucose uptake in fibroblasts. Bioluminescence-based glucose uptake analysis using the Glucose Uptake-Glo™ Assay shows that the amount of glucose taken up by fibroblasts (n = 2) is moderately increased under compression (*p* = 0.03, though this experiment was not sufficiently powered). Luminescence counts were normalized to cell count per well. Error bars represent SEM.

**Figure S3.**
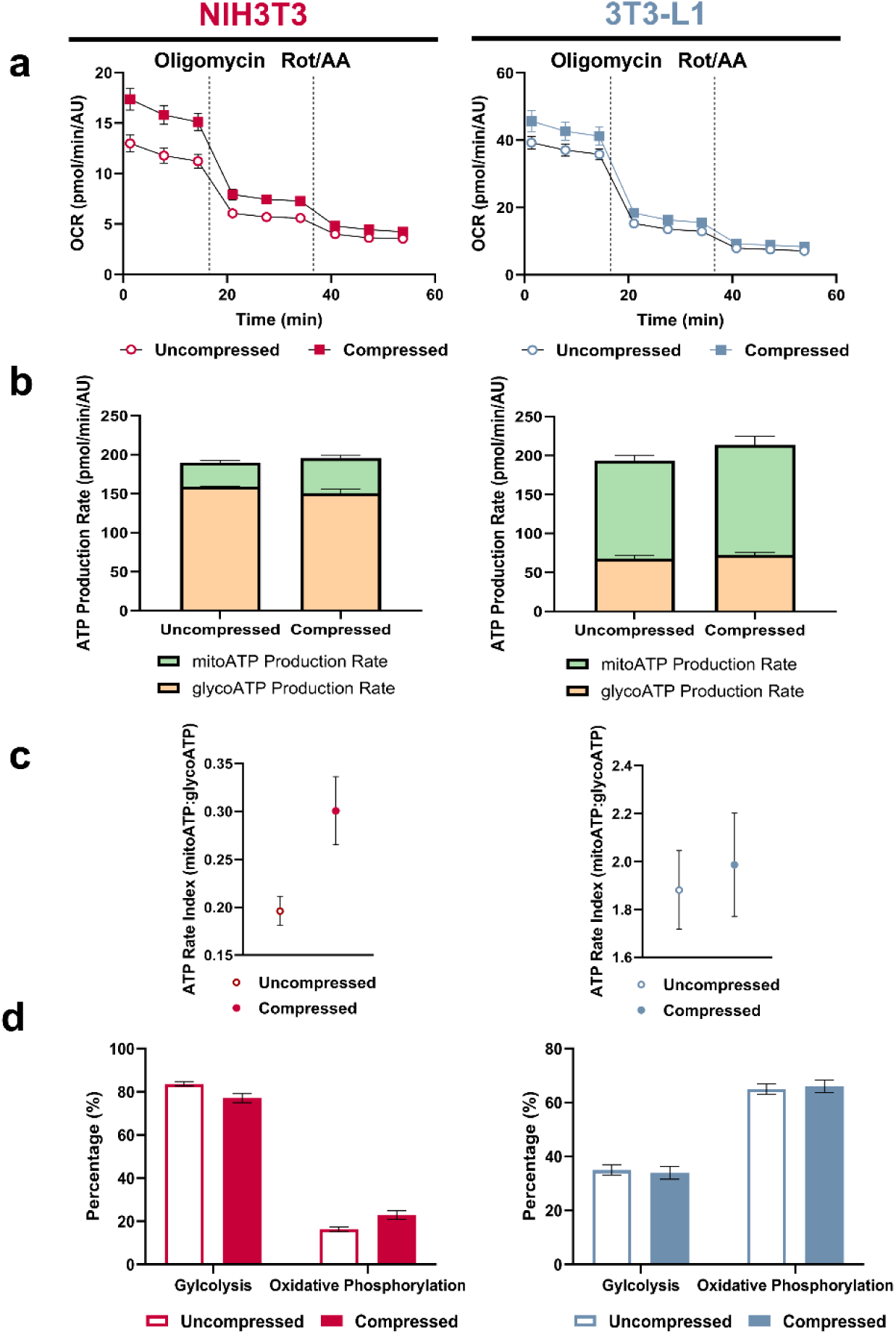
Relative glycolytic and mitochondrial ATP production does not vary between compressed and uncompressed stromal cells as assessed by Seahorse assay. **a)** Representative plots of oxygen consumption rate (OCR) over time for NIH3T3 (left; n = 3) and 3T3-L1 (right; n = 3) cells. **b)** Relative contribution to ATP production through glycolysis (glycoATP) and oxidative phosphorylation (mitoATP). **c)** ATP rate index (proportion of mitoATP:glycoATP) for compressed and uncompressed cells. Higher values indicate a preference for energy production via oxidative phosphorylation while lower values suggest a bias towards glycolysis-driven ATP production. **d)** Percent activity of glycolysis and oxidative phosphorylation is not significantly different between compressed and uncompressed cells. When considered alongside our FLIM results in Fig. 2, these findings suggest that FLIM offers superior sensitivity in detecting *in situ* metabolic changes in our system compared to traditional metabolic assays such as Seahorse. Error bars represent SEM and asterisks indicate statistical significance (\**p* < 0.05, \*\**p* < 0.01, \*\*\**p* < 0.001, \*\*\*\**p* < 0.0001) determined using a Student’s t-test.

**Figure S4.**
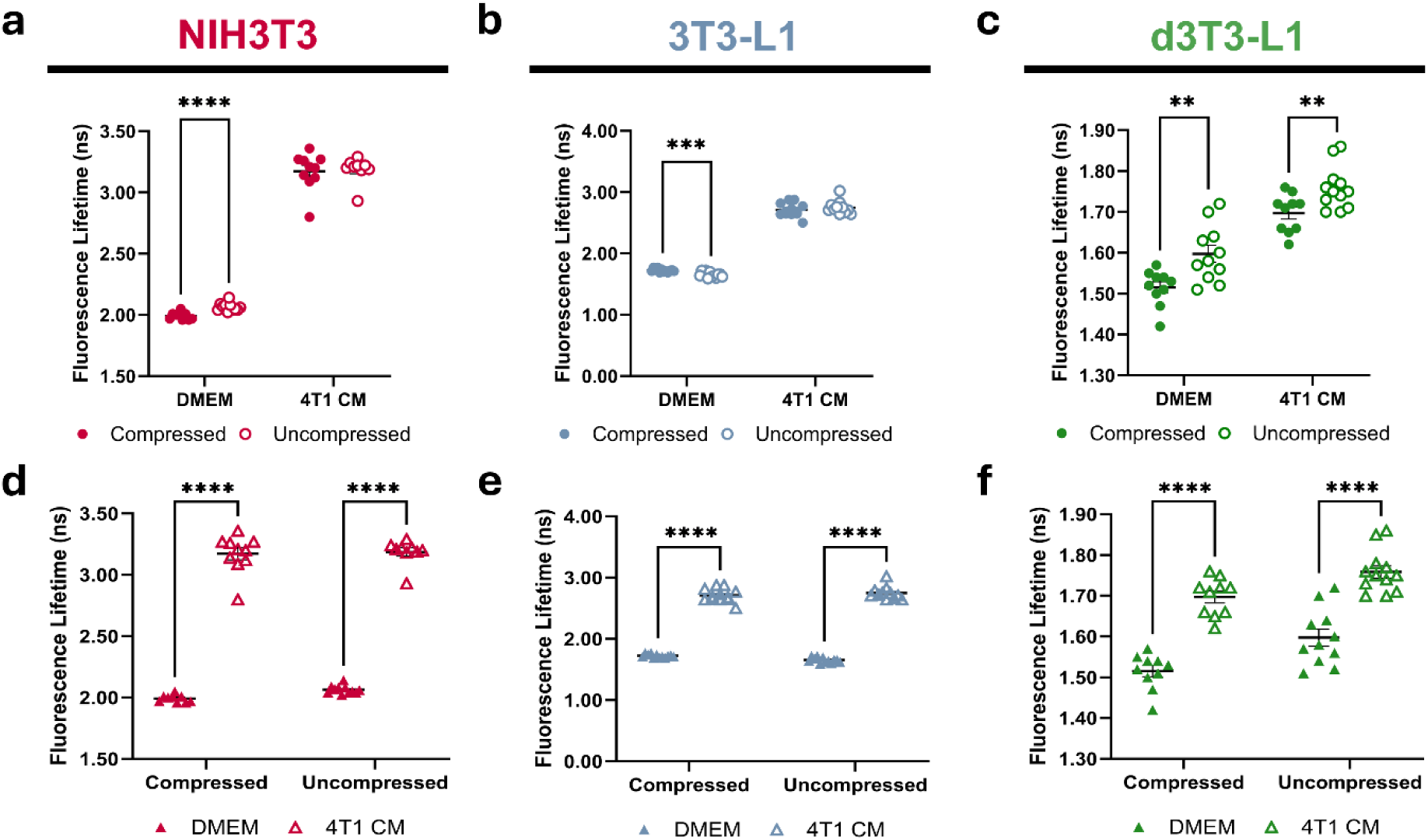
Comparison of fluorescence lifetime between compressed and uncompressed stromal cells cultured in DMEM or 4T1-conditioned media (CM). The inclusion of 4T1 CM counteracts compression-induced metabolic reprogramming in **a)** NIH3T3 (data from Fig. 2d and Fig. 6d) and **b)** 3T3-L1 cells (data from Fig. 2e and Fig. 6e), but not in **c)** d3T3-L1 cells (data from Fig. 2f and Fig. 6f). Additionally, 4T1 CM significantly increases fluorescence lifetime regardless of compression status in **d)** NIH3T3 (data from Fig. 2d and Fig. 6d), **e)** 3T3-L1 (data from Fig. 2e and Fig. 6e), and **f)** d3T3-L1 (data from Fig. 2f and Fig. 6f) cells. Error bars represent SEM. Statistical significance was determined using a Student’s t-test, with asterisks indicating significance levels (\**p* < 0.05, \*\**p* < 0.01, \*\*\**p* < 0.001, \*\*\*\**p* < 0.0001).

