## Supplemental Table 1 for "Instant fluorescence lifetime imaging microscopy reveals mechano-metabolic reprogramming of stromal cells in breast cancer peritumoral microenvironments"

**Supplemental Table 1. Differential expression of transcripts involved in glycolysis and oxidative phosphorylation for compressed vs. uncompressed NIH3T3 fibroblasts**

| **Gene Name** | **log2FC** | **padj** |
| --- | --- | --- |
| **Glycolysis^1^** | | |
| Slc4a4 | 0.600 | 1.96E-02 |
| Ppargc1a | 0.582 | 5.01E-02 |
| Htr2a | 0.466 | 6.35E-02 |
| Ier3 | 0.454 | 2.58E-01 |
| Insr | 0.376 | 2.77E-03 |
| Stat3 | 0.353 | 1.18E-03 |
| Pfkfb2 | 0.320 | 2.10E-01 |
| Pfkfb3 | 0.289 | 2.04E-01 |
| Ep300 | 0.275 | 2.99E-02 |
| Igf1 | 0.259 | 3.81E-01 |
| Lipa | 0.240 | 5.90E-02 |
| Psen1 | 0.206 | 6.51E-02 |
| Ncor1 | 0.190 | 1.30E-01 |
| Sik2 | 0.179 | 2.71E-01 |
| Foxk1 | 0.175 | 1.86E-01 |
| Ogt | 0.158 | 1.17E-01 |
| Hk1 | 0.147 | 1.70E-01 |
| App | 0.141 | 1.44E-01 |
| Mfsd8 | 0.139 | 5.80E-01 |
| Kat2b | 0.135 | 3.06E-01 |
| Pfkm | 0.125 | 2.39E-01 |
| Hk2 | 0.123 | 3.37E-01 |
| Hif1a | 0.110 | 2.97E-01 |
| Zbtb7a | 0.104 | 3.48E-01 |
| Galt | 0.093 | 6.69E-01 |
| Bcl2l13 | 0.082 | 4.80E-01 |
| Jmjd8 | 0.074 | 8.11E-01 |
| Mpi | 0.073 | 6.43E-01 |
| Bpgm | 0.063 | 7.10E-01 |
| Eno2 | 0.039 | 8.27E-01 |
| Src | 0.038 | 7.78E-01 |
| Myc | 0.017 | 9.29E-01 |
| Prkag2 | 0.017 | 9.36E-01 |
| Eno3 | 0.016 | 9.44E-01 |
| Trex1 | 0.009 | 9.79E-01 |
| Ddit4 | 0.008 | 9.68E-01 |
| Flcn | 0.003 | 9.88E-01 |
| Nupr1 | 0.002 | 9.92E-01 |
| Pgam1 | -0.009 | 9.44E-01 |
| Prkaca | -0.015 | 9.25E-01 |
| Aldoart1 | -0.016 | 9.70E-01 |
| Ucp2 | -0.023 | 8.73E-01 |
| Prkag1 | -0.034 | 8.06E-01 |
| Mlx | -0.035 | 8.29E-01 |
| Pfkp | -0.037 | 8.34E-01 |
| Tkfc | -0.039 | 8.37E-01 |
| Prkaa1 | -0.040 | 7.37E-01 |
| Enpp1 | -0.054 | 7.55E-01 |
| Pfkl | -0.060 | 5.84E-01 |
| Gpi1 | -0.066 | 5.14E-01 |
| Col6a1 | -0.079 | 4.25E-01 |
| Sirt6 | -0.080 | 6.04E-01 |
| Aldoa | -0.089 | 3.53E-01 |
| Khk | -0.100 | 8.38E-01 |
| Eno4 | -0.111 | 8.22E-01 |
| Pfkfb1 | -0.121 | 6.97E-01 |
| Fkrp | -0.121 | 3.28E-01 |
| Tigar | -0.133 | 4.95E-01 |
| Arl2 | -0.136 | 2.46E-01 |
| Mtch2 | -0.136 | 1.46E-01 |
| Pkm | -0.139 | 1.05E-01 |
| Pgk1 | -0.147 | 7.67E-02 |
| Slc2a6 | -0.164 | 6.94E-01 |
| Prxl2c | -0.178 | 1.27E-01 |
| Eno1 | -0.204 | 2.35E-02 |
| Galk1 | -0.215 | 1.15E-01 |
| Gapdh | -0.225 | 1.58E-02 |
| Eno1b | -0.232 | 2.19E-02 |
| Ppp2ca | -0.232 | 4.15E-02 |
| Tpi1 | -0.303 | 5.77E-03 |
| Gale | -0.538 | 4.33E-05 |
| **Oxidative Phosphorylation^2^** | | |
| Atp7a | 0.521 | 8.36E-03 |
| Pink1 | 0.350 | 1.87E-02 |
| Slc25a23 | 0.143 | 6.95E-01 |
| Tmem135 | 0.131 | 5.01E-01 |
| Bcl2l13 | 0.082 | 4.80E-01 |
| Afg1l | 0.053 | 8.48E-01 |
| Sdhc | 0.034 | 7.91E-01 |
| Iscu | 0.030 | 8.56E-01 |
| Myc | 0.017 | 9.29E-01 |
| Bid | 0.013 | 9.53E-01 |
| Nipsnap2 | 0.008 | 9.67E-01 |
| Nupr1 | 0.002 | 9.92E-01 |
| Coq9 | -0.005 | 9.81E-01 |
| Slc25a33 | -0.006 | 9.76E-01 |
| Dguok | -0.007 | 9.78E-01 |
| Ndufs1 | -0.009 | 9.53E-01 |
| Sdha | -0.020 | 8.71E-01 |
| Msh2 | -0.029 | 8.39E-01 |
| Ndufv1 | -0.029 | 8.26E-01 |
| Macroh2a1 | -0.030 | 8.11E-01 |
| Bdnf | -0.036 | 7.77E-01 |
| Ndufaf1 | -0.043 | 8.63E-01 |
| Dld | -0.051 | 6.50E-01 |
| Antkmt | -0.056 | 7.56E-01 |
| Tafazzin | -0.059 | 7.01E-01 |
| Ndufv3 | -0.065 | 6.75E-01 |
| Vcp | -0.077 | 4.36E-01 |
| Cox6a2 | -0.082 | 8.32E-01 |
| 1700066M21Rik | -0.085 | 6.36E-01 |
| Ndufa10 | -0.087 | 4.05E-01 |
| Uqcrc1 | -0.091 | 5.00E-01 |
| Sdhd | -0.092 | 3.94E-01 |
| Ndufs3 | -0.107 | 3.71E-01 |
| Atp5f1a | -0.115 | 1.86E-01 |
| Cox5a | -0.121 | 2.23E-01 |
| Ndufs7 | -0.126 | 3.37E-01 |
| Coq7 | -0.135 | 3.43E-01 |
| Mtch2 | -0.136 | 1.46E-01 |
| Uqcrfs1 | -0.148 | 2.03E-01 |
| Sdhaf2 | -0.149 | 2.09E-01 |
| Park7 | -0.152 | 2.55E-01 |
| Uqcc3 | -0.152 | 3.93E-01 |
| Atp5po | -0.153 | 2.13E-01 |
| Atp5f1b | -0.159 | 1.02E-01 |
| Ccnb1 | -0.162 | 1.19E-01 |
| Ndufs2 | -0.164 | 2.04E-01 |
| Cyc1 | -0.166 | 9.14E-02 |
| Atpsckmt | -0.167 | 4.06E-01 |
| Shmt2 | -0.172 | 7.44E-02 |
| Atp5pb | -0.180 | 1.09E-01 |
| Atp5f1d | -0.180 | 8.28E-02 |
| Cox6a1 | -0.182 | 1.58E-01 |
| Cox5b | -0.182 | 7.43E-02 |
| Tefm | -0.200 | 3.36E-01 |
| Ndufs6 | -0.211 | 1.30E-01 |
| Cox4i1 | -0.216 | 7.41E-02 |
| Ndufs8 | -0.232 | 1.87E-02 |
| Atp5pf | -0.232 | 3.53E-02 |
| Atp5f1c | -0.235 | 4.70E-02 |
| Ndufv2 | -0.247 | 8.28E-02 |
| Ndufa7 | -0.248 | 4.75E-02 |
| Atp5f1e | -0.282 | 2.72E-02 |
| Ndufb6 | -0.282 | 6.79E-02 |
| Uqcr10 | -0.283 | 4.44E-02 |
| Uqcc2 | -0.289 | 2.92E-02 |
| Cdk1 | -0.292 | 1.30E-02 |
| Dnajc15 | -0.295 | 1.81E-02 |
| Fxn | -0.295 | 6.38E-02 |
| Uqcrq | -0.301 | 2.81E-02 |
| Atp5pd | -0.319 | 1.13E-02 |
| Coa6 | -0.324 | 6.95E-02 |
| Atp5me | -0.362 | 7.50E-03 |
| Uqcrh | -0.389 | 5.41E-03 |
| Ndufa12 | -0.400 | 3.33E-03 |
| Cox7c | -0.401 | 2.78E-03 |
| Cycs | -0.409 | 1.95E-03 |
| Uqcrb | -0.416 | 5.13E-03 |
| Atp5mf | -0.506 | 5.57E-02 |
| Cox7a2 | -0.532 | 2.95E-04 |

^1^Blue = transcripts involved in glycolysis

^2^Red = transcripts involved in oxidative phosphorylation
