## Supplemental Table 2 for "Instant fluorescence lifetime imaging microscopy reveals mechano-metabolic reprogramming of stromal cells in breast cancer peritumoral microenvironments"

**Supplemental Table 2. List of top differentially expressed genes for compressed vs. uncompressed NIH3T3 fibroblasts**

| **Gene Name** | **log2FC** | **padj** |
| --- | --- | --- |
| C3 | 3.79 | 5.05E-06 |
| Cxcl5 | 2.56 | 3.37E-06 |
| Serpina3h | 2.45 | 2.75E-11 |
| Cxcl1 | 1.97 | 5.22E-04 |
| Chi3l1 | 1.71 | 1.13E-05 |
| Ptprv | 1.36 | 4.88E-05 |
| Klhl30 | 1.29 | 1.69E-04 |
| Siglecg | 1.23 | 2.79E-05 |
| Tnfaip3 | 1.15 | 3.89E-05 |
| C1s1 | 1.11 | 2.53E-03 |
| Fas | 1.10 | 8.13E-05 |
| Glipr1 | 1.04 | 2.36E-03 |
| Crispld2 | 1.04 | 8.40E-03 |
| Col5a3 | 1.04 | 8.23E-04 |
| Slpi | 1.03 | 1.50E-06 |
| Mmp9 | 1.02 | 2.86E-04 |
